# Molecular subtyping based on hippocampal cryptic exon burden reveals proteome-wide changes associated with TDP-43 pathology across the spectrum of LATE and Alzheimer’s Disease

**DOI:** 10.1101/2025.05.30.656396

**Authors:** Adam N. Trautwig, Anantharaman Shantaraman, Mingee Chung, Eric B. Dammer, Lingyan Ping, Duc M. Duong, Leonard Petrucelli, Michael E. Ward, Jonathan D. Glass, Peter T. Nelson, Allan I. Levey, Zachary T. McEachin, Nicholas T. Seyfried

## Abstract

TDP-43 pathology is a defining feature of Limbic-Predominant Age-Related TDP-43 Encephalopathy neuropathologic change (LATE-NC) and is frequently comorbid with Alzheimer’s disease neuropathologic change (ADNC). However, the molecular consequences of co-occurring LATE-NC and ADNC pathology (TDP-43, β-amyloid, and tau protein pathologies) remain unclear. Here, we conducted a comparative biochemical, molecular, and proteomic analysis of hippocampal tissue from 90 individuals spanning control, LATE-NC, ADNC, and ADNC+LATE-NC groups to assess the impact of cryptic exon (CE) inclusion, phosphorylated TDP-43 pathology (pTDP-43), and AD-related pathologies (β-amyloid, and tau) on the proteome. ADNC+LATE-NC cases exhibited the highest burden of CE inclusion as quantified by measuring the levels of known TDP-43 regulated CEs within eight transcripts: *STMN2, UNC13A, ELAVL3, KALRN, ARHGAP32, CAMK2B, PFKP,* and *SYT7*. While CE levels correlated with pTDP-43 pathology, they were more strongly correlated with each other, suggesting that the molecular signature of CE inclusion may serve as a more sensitive measure of TDP-43 dysfunction than pTDP-43 pathology alone. Unbiased classification based on the relative abundance of these eight CEs stratified individual cases into low, intermediate, and high CE burden subtypes, largely independent of β-amyloid and tau pathology. Proteome-wide correlation analysis revealed a bias toward reduced protein levels from genes harboring TDP-43-regulated CEs in cases with high cumulative CE burden. Notably, proteins significantly decreased under high CE burden included canonical STMN2, ELAVL3, and KALRN, as well as kinesin proteins that are genetically associated with amyotrophic lateral sclerosis. Co-expression network analysis identified both shared and distinct biological processes across CE subtypes and pathways associated with pTDP-43, tau, β-amyloid pathologies, and CE accumulation in the hippocampus. Protein modules associated with TDP-43 loss of function were prioritized by integrating proteomic data from TDP-43–depleted human neurons with the hippocampal co-expression network. Specifically, we observed decreased endosomal vesicle, microtubule-binding, and synaptic modules, alongside an increase in RNA-binding modules. These results provide new insights into the proteomic impact of CE burden across the spectrum of LATE and AD pathological severity, highlighting the molecular consequences of TDP-43 dysfunction in neurodegenerative disease

## Introduction

Alzheimer’s disease neuropathologic change (ADNC) is defined by two hallmark brain pathologies: β-amyloid (Aβ) plaques and tau neurofibrillary tangles (NFTs)^1^. However, the large majority of brains with ADNC and documented dementia harbor one or more additional age-related pathologies at autopsy including but not limited to cerebrovascular disease, Lewy body inclusions, and TAR DNA-binding protein 43 (TDP-43) aggregates^2-5^. TDP-43 pathology, characterized by aberrant phosphorylation and mislocalization as detected by immunohistochemistry, is observed in approximately half of pathologically confirmed ADNC dementia case^6, 7^. However, TDP-43 pathology is not a defining feature of ADNC itself. In the context of aging-related dementia, the presence of TDP-43 pathology is a hallmark of limbic-predominant age-related TDP-43 encephalopathy neuropathologic change (LATE-NC), which typically begins in the amygdala, progresses to the hippocampus, and eventually affects the middle frontal gyrus^8, 9^. Across a broad range of cognitive conditions, LATE-NC was observed in approximately one-third of autopsied individuals past age 85 in large community-based autopsy studies^8, 10, 11^. LATE-NC can occur in the absence or presence of ADNC and is associated with significant clinical impairment independent of other pathologies^8, 12^. Although it is known that cases with comorbid ADNC and LATE-NC tend to suffer a relatively severe clinical course, the mechanisms by which TDP-43, Aβ and tau pathologies interact to accelerate cognitive decline remain unknown.

TDP-43 was initially identified as a neuropathological hallmark in the context of amyotrophic lateral sclerosis (ALS) and a subtype of frontotemporal lobar dementia (FTLD-TDP)^13^. Since the initial discovery that TDP-43 forms phosphorylated (pTDP-43), intracellular aggregates in neurodegenerative diseases, several studies have demonstrated that TDP-43 aggregates can promote toxicity through toxic gain-of-function mechanisms^14-16^. However, the depletion or loss of nuclear TDP-43 is also considered a hallmark of TDP-43 proteinopathies^17^ and can occur pre-symptomatically^18, 19^, preceding the formation of cytosolic aggregates, and suggesting that TDP-43 loss of function is an early event in disease pathogenesis. As an RNA-binding protein, TDP-43 has well-established roles in alternative splicing and mRNA stability^20, 21^ and is crucial for maintaining normal levels and splicing patterns of thousands of mRNAs, many of which are mis-spliced in TDP-43 proteinopathies due to its dysfunction^22^. High-throughput sequencing, combined with cross-linking immunoprecipitation experiments have identified direct TDP-43 mRNA targets and revealed that most of its binding sites are in “UG”-rich dinucleotide repeats within introns of pre-mRNAs^21, 22^. Many of these intronic binding sites were later found to function as splicing repressor sites for TDP-43^23^. Thus, a key consequence of TDP-43 loss or dysfunction is the erroneous inclusion of non-conserved intronic sequences in mature RNA transcripts, known as cryptic exons (CE)^24, 25^.

*In silico* prediction of CE sequences suggest that these novel RNA transcripts are substrates for RNA surveillance mechanisms that result in reduced expression^24, 26^. Consequently, cryptic exons are anomalously included in critical transcripts such as *STMN2* and *UNC13A* which can produce truncated or destabilized RNAs and lead to a loss of expression through nonsense-mediated decay (NMD) or pre-mature polyadenylation ^27-31^. Thus, the presence of CEs essentially serves as a surrogate marker for TDP-43 deficiency in cells and tissues. However, the inclusion of CEs in certain genes regulated by TDP-43 can lead to the translation of novel protein isoforms^19, 32^. Some of these *de novo* proteins, like HDGFL2, have emerged as biomarkers of TDP-43 dysfunction in biofluids and tissues^19, 32^. To date, the global impact of TDP-43 dysfunction and resulting cryptic exon inclusion on the proteome remains unknown.

A recent study from our group investigated the accumulation of eight TDP-43-regulated CEs in the hippocampus of LATE-NC cases and found several CEs strongly expressed, regardless of ADNC co-pathology^33^. Here, we expanded these studies by conducting comparative proteomics on 90 hippocampal tissues spanning the clinicopathological spectrum of control, LATE-NC, ADNC, and ADNC+LATE-NC cases to assess the impact of pTDP-43, tau, and amyloid pathologies on the proteome. We also sought to better understand the relationship with AD pathology and TDP-43 pathologies including the impact of CE inclusion. Given that TDP-43 mis-localization occurs early and potentially precedes pathological TDP-43 aggregation, we hypothesized that CE inclusion (loss of function) and pTDP-43 levels (gain of function) may not always co-occur and could represent distinct stages or mechanisms of disease progression. To explore this, we used correlation analyses among the CEs measured and distinct neuropathologies. Furthermore, we used an unbiased classification approach based on the relative abundance of eight CE-containing transcripts to stratify hippocampal cases into subtypes with low, intermediate, or high CE burden, that were largely independent of β-amyloid and tau pathology. We then examined how CE burden relates to proteomic alterations in the brain by performing proteome-wide correlation and co-expression network analyses. Because CE inclusion is predicted to reduce expression of full-length proteins via NMD or pre-mature polyadenylation, we specifically assessed whether proteins encoded by CE-containing transcripts such as canonical *STMN2*, *ELAVL3*, and *KALRN* were reduced in high CE burden cases. To validate and extend these findings, we integrated postmortem brain proteomic data with proteomes derived from TDP-43-depleted human iPSC-derived neurons, aiming to identify conserved molecular pathways and compensatory mechanisms associated with CE burden and TDP-43 dysfunction. Through this approach, we aimed to uncover the proteomic signatures and potential biomarkers linked to CE inclusion, which define the broader impact of TDP-43 loss of function in human neurodegenerative disease.

## Materials and Methods

### Case Selection

Human post-mortem tissue from the hippocampi of 91 individuals (**Supplemental Table 1**) was selected from the University of Kentucky AD Research Center (ADRC) autopsy community-based cohort; details of the cohort, inclusion/exclusion criteria, etc., and clinical diagnostic workups were previously described^10, 34^. For the purposes of the present study, a convenience sample was selected of hippocampi snap-frozen using liquid nitrogen at the time of autopsy (with post-mortem interval <4hrs) and stored in -80°C freezers thereafter. In terms of neuropathologic diagnostic assessments, stage-based diagnoses of LATE-NC, ADNC, and other pathologies used consensus-based criteria^10, 35-37^. None of the included participants had FTD/FTLD, ALS, or other rare (<1:300 lifetime risk) conditions.

### Isobaric Tandem Mass Tag (TMT) Peptide Labeling

Tissue homogenization and protein (e.g. trypsin and LysC) digestion of all cases was performed essentially as described^38, 39^. Prior to labeling, these 91 cases were randomized into 6 batches by age, sex, and diagnosis using ARTS (automated randomization of multiple traits for study design)^40^. Labeling was performed using TMTpro 18-plex kits (ThermoFisher 44520). Each batch included two TMT channels with a labeled GIS standard. Labeling of sample peptides was performed as previously described^38, 41, 42^. Briefly, each sample (100 μg of peptides) was re-suspended in 100 mM triethylammonium bicarbonate (TEAB) buffer (100 μL). TMT labeling reagents (5 mg) were equilibrated to room temperature. Anhydrous ACN (256 μL) was added to each reagent channel. Each channel was then gently vortexed for 5 minutes. A volume of 41 μL from each TMT channel was transferred to each peptide solution and allowed to incubate for 1 hour at room temperature. The reaction was quenched with 5% (vol/vol) hydroxylamine (8 μL) (Pierce). All channels were then dried by SpeedVac (LabConco) to approximately 150 μL, diluted with 1 mL of 0.1% (vol/vol) TFA, and acidified to a final concentration of 1% (vol/vol) FA and 0.1% (vol/vol) TFA. Labeled peptides were desalted with a 200 mg C18 Sep-Pak column (Waters). Prior to sample loading, each Sep-Pak column was activated with 3 mL of methanol, washed with 3 mL of 50% (vol/vol) ACN, and equilibrated with 2×3 mL of 0.1% TFA. After sample loading, each column was washed with 2×3 mL 0.1% (vol/vol) TFA followed by 2 mUPL of 1% (vol/vol) FA. Elution was performed with 2 volumes of 1.5 mL 50% (vol/vol) ACN. The eluates were then dried to completeness by SpeedVac. Subsequent high pH fractionation of all cases was performed essentially as described ^38, 43^ A total of 192 individual equal volume fractions were collected across the gradient and subsequently pooled by concatenation into 96 fractions^43^. The fractions were then dried to completeness using a SpeedVac.

### TMT Mass Spectrometry (TMT-MS) Analysis

MS analysis was performed on the high-pH fractionated samples as previously described with modifications^38, 39, 44, 45^. Briefly, fractions were resuspended in an equal volume of loading buffer (0.1% FA, 0.03% TFA, 1% ACN) and analyzed by liquid chromatography coupled to tandem mass spectrometry (LC-MS/MS). Peptide eluents were separated on a custom in-house packed Charged Surface Hybrid (CSH) column (1.7 um, 15 cm × 150 μM internal diameter) by a Dionex RSLCnano ultra-performance liquid chromatography (UPLC) system (ThermoFisher Scientific). Buffer A comprised water with 0.1% (vol/vol) FA, and buffer B comprised 80% (vol/vol) ACN in water with 0.1% (vol/vol) FA. Elution was performed over a 30 min gradient with a flow rate of 1500 nL/min. The gradient ranged from 1% to 99% solvent B. Peptides were monitored on a Orbitrap Eclipse mass spectrometer with high-field asymmetric waveform ion mobility spectrometry (FAIMS) (FAIMS Pro Interface, ThermoFisher Scientific). Two compensation voltages were chosen for FAIMS. For each voltage (-45 and -65) top speed cycle of 1.5 seconds, the full scan (MS1) was performed with an m/z range of 410-1600 and 60,000 resolution at standard settings. The higher energy collision-induced dissociation (HCD) tandem scans were collected at 35% collision energy with an isolation of 0.7 m/z, resolution of 30,000 with TurboTMT, AGC setting of 250% normalized AGC target, and a maximum injection time of 54 ms. For all batches, dynamic exclusion was set to exclude previously sequenced peaks for 20 seconds within a 10-ppm isolation window.

### Database Search and Protein Quantification for TMT-MS Hippocampus Proteome

Database searches and protein quantification was performed on 576 RAW files (96 RAW files/fractions per batch) using FragPipe (version 20.0). The FragPipe pipeline relies on MSFragger (version 4.0)^46, 47^ for peptide identification, MSBooster^48^, and Percolator^49^ for FDR filtering and downstream processing. MS/MS spectra were searched against all canonical Human proteins downloaded from Uniprot (20,402; accessed 02/11/2019), as well as common contaminants (51 total), and all reverse sequences (20,453). The workflow we used in FragPipe followed default TMT-18 plex (i.e., TMTpro) parameters. Briefly, precursor mass tolerance was -20 to 20 ppm, fragment mass tolerance of 20 ppm, mass calibration and parameter optimization were selected, and isotope error was set to -1/0/1/2/3. Enzyme specificity was set to strict-trypsin with up to two missing trypsin cleavages allowed. Cleavage was set to fully tryptic. Peptide length was allowed to range from 7 to 35 and peptide mass from either 200 to 5,000 Da. Variable modifications that were allowed in our search included: oxidation on methionine, N-terminal acetylation on protein and peptide, and phosphorylation on S, T, and Y with a maximum of 3 variable modifications per peptide^50^. The false discovery rate (FDR) threshold was set to 1%. A total of 191,342 peptides which mapped to 10,746 proteins were detected and column normalized (**Supplemental Table 2**). After filtering out proteins that were absent in 50% or more of specimens 108,842 peptides and 9,537 proteins were retained.

### Bioinformatics Processing and Statistical Analysis

We employed a Tunable Approach for Median Polish of Ratio (TAMPOR)^51^ and removal of peptides or proteins absent in 50% of cases or greater, as previously published^52, 53^. To ensure the reliability of our data, we initiated the analysis by identifying and removing potential outliers, initially one outlier AD case was removed in two iterations resulting in the final dataset of (n=90 cases). We then performed parallelized^54^ ordinary, nonparametric, bootstrapping regression to remove variation due to age, sex, post-mortem interval (PMI), and batch effect using an established pipeline (**Supplemental Table 3)**^44, 45^. Fast parallel one-way ANOVA with Benjamini-Hochberg correction for multiple comparisons was conducted within each disease group using an in-house script (https://github.com/edammer/parANOVA) to identify peptides and proteins that were differentially abundant (**Supplemental Table 4**). Differential abundance is presented as volcano plots, which were generated with the ggplot2 package^55^. Correlations were performed using the biweight midcorrelation function as implemented in the WGCNA R package, using only pairwise complete observations for the calculations (**Supplemental Table 5)**. Phosphorylated TDP-43 was imputed with minimum values equivalent to 0.1, or roughly half of the lowest detected level.

### Molecular Subtyping Based on CE Profiles

Relative expression of CEs were determined using qPCR, as previously described^33^, whereby relative expression was normalized to RPLP0, GAPDH, and CYC1. Briefly, the geometric mean of normalization protein Ct values was subtracted from each cryptic exon (ΔCt) and average control group ΔCt were subtracted from each ΔCt (*i.e.* ΔΔCt). Relative expression was log2 transformed for downstream analyses. Individual abundances across 8 CEs (STMN2, UNC13A, ELAVL3, KALRN, ARHGAP32, CAMK2B, PFKP, and SYT7) were hierarchically clustered, unsupervised and then binned based on clustering dendrogram and a visual inspection of expression pattern. Supervised binning of unsupervised hierarchical clustering was validated using principal component analysis (PCA) as implemented in the prcomp function of the stats R package. We used two publicly available data sources (**Supplemental Table 6)**^30, 56^ in order to compile a list of 183 CE transcripts. Proteins in this list were highlighted in scatterplots and volcano plots to visualize overarching trends in correlation (to burden index) and abundance in TDP-43 KD iNeurons. The Spearman rank correlation was performed on the cumulative cryptic exon (CE) rank for each of the eight CE transcripts (**Supplemental Table 7)**. For each subject (n=90), CE abundance was ranked from 1 (lowest) to 90 (highest) and summed across all eight CEs to generate a Cryptic Exon Burden Score per individual case.

### Protein Network Analysis

Weighted Gene Co-expression network analysis (WGCNA)^57^ was used to construct modules of co-expressing proteins as previously published^52, 53^. Briefly, the blockwiseModules function from the WGCNA package in R was utilized with the following parameters: soft threshold power beta = 24.5, deepSplit = 2, minimum module size = 15, merge cut height = 0.07, and a signed network with partitioning around medoids (mapping a distance matrix to k clusters, where K is data-adaptively selected; **Supplemental Table 8**)^58^. Module correlation to disease type was evaluated with biweight midcorrelation (BiCor) analysis by separating each disease control combination. Fisher’s exact test (FET) was performed for each module’s members against the merged human brain cell type marker list to determine cell type enrichment using an in-house script (https://github.com/edammer/cellTypeFET). Similarly, to determine module gene ontology (GO) a FET was performed for each module member against the Bader Lab’s GMT formatted ontology lists from February 8, 2023^59^ (https://github.com/edammer/GOparallel; **Supplemental Table 9**). One way ANOVA was conducted across all molecular subtypes for each module eigenprotein.

### CRISPRi Knockdown of TDP-43 and Differentiation into Neurons

CRISPRi knockdown experiments were performed using the i11w-mNC line, a derivative of the previously described WTC11 line^32^. This line stably expresses CAG-dCas9-BFP-KRAB at the CLYBL safe harbor locus and a tetracycline-inducible mNGN2 transgene at the AAVS1 locus. CRISPRi knockdown (KD) of TDP-43 was achieved by infecting iPSCs with lentiviruses delivering sgRNAs targeting TARDBP or a non-targeting control sgRNA. Neurons were differentiated as previously described^32^ and harvested 17 days post-differentiation for all experiments. Both control and TDP-43 KD lines were generated in biological quadruplicate (n=4 per group).

### Data Independent Acquisition (DIA-MS) Mass Spectrometry of Human Neurons

Protein lysates (20 ug) from the iNeuron cell lines were digested with trypsin and endopeptidase LysC as previously described^53^. Each sample was resuspended in 25 uL of loading buffer (0.1% FA) and 0.5 ul was analyzed by liquid chromatography coupled to tandem mass spectrometry. Peptide eluents were separated on uPAC high throughput column by a Vanquish Neo (ThermoFisher Scientific). Buffer A was water with 0.1% (vol/vol) formic acid, and buffer B was 100% (vol/vol) acetonitrile in water with 0.1% (vol/vol) formic acid. Elution was performed over a 5 min (short) or 30 min (long) elution gradient. The gradient was from 1% to 90% solvent B. Peptides were monitored by DIA on a Orbitrap Astral mass spectrometer (ThermoFisher Scientific) fitted with a high-field asymmetric waveform ion mobility spectrometry (FAIMS Pro) ion mobility source (ThermoFisher Scientific). One compensation voltages (CV) of -35 was chosen for the FAIMS. Each cycle consisted of one full scan (MS1) was performed with an m/z range of 380-980 at 240,000 resolution, 500% AGC and 3 ms injection time. The higher energy collision-induced dissociation (HCD) DIA scans were collected with a 2 m/z isolation windows over the entire precursor range (380-980 m/z) with a time of 0.6 seconds and 2.5 ms injection time. Collision energy was set to 25% and scan range set to 150 - 2000 m/z. Samples for the 5 min gradient were collected in technical replicate (n=8 DIA-MS runs per group). Library-free database searches and protein quantification were performed on the DIA-MS raw files using Spectronaut (version 18.1) with default fully tryptic parameter settings. The search database was identical to that used for the TMT-MS analysis, but also included cryptic peptide sequences from previously predicted de novo proteins. Log-transformed MS2-based quantification was used to assess relative abundance. For the 5-minute gradient, data from each technical replicate was averaged per sample. Missing values for both the 5-minute and 30-minute gradients were imputed using Perseus-style imputation (**Supplemental Table 10-11**). T-Test were performed against for each protein/peptide detected against iNeuron type (Control or TDP-43 KD; **Supplemental Table 12-13**). The DIA-MS raw files, protein database and Spectronaut output are available on Synapse.

## Results

### Pathological, biochemical, and molecular characterization of LATE with or without AD pathology

To investigate proteome-wide differences associated with Alzheimer’s disease (AD) and TDP-43 pathology, we selected three groups of cases: those with predominantly TDP-43 neuropathological changes (referred to as LATE-NC, or simply LATE), those with only Aβ plaque and tau tangle neuropathological changes (ADNC, or simply AD), and those with both AD and TDP-43 neuropathology (ADNC+LATE-NC or simply AD+LATE). We performed TMT-MS-based proteomic analysis (**Figure 1A**) on 91 postmortem hippocampal tissue from individuals classified into four groups: controls (n = 21), AD (n = 25), AD+LATE (n = 35), and LATE (n = 10). The hippocampus was selected for analysis due to its heightened vulnerability to co-occurring pathologies in AD and related dementias including LATE^8, 12^. Case characteristics are detailed in **Supplemental Table 1**. Following quality control and outlier removal, our proteomic analysis identified 9,537 proteins across 90 cases with one AD case removed as an outlier. Protein abundance was adjusted for batch effects, postmortem interval (PMI), age, and sex across groups (**Supplemental Figure 1**).

**Figure 1.**
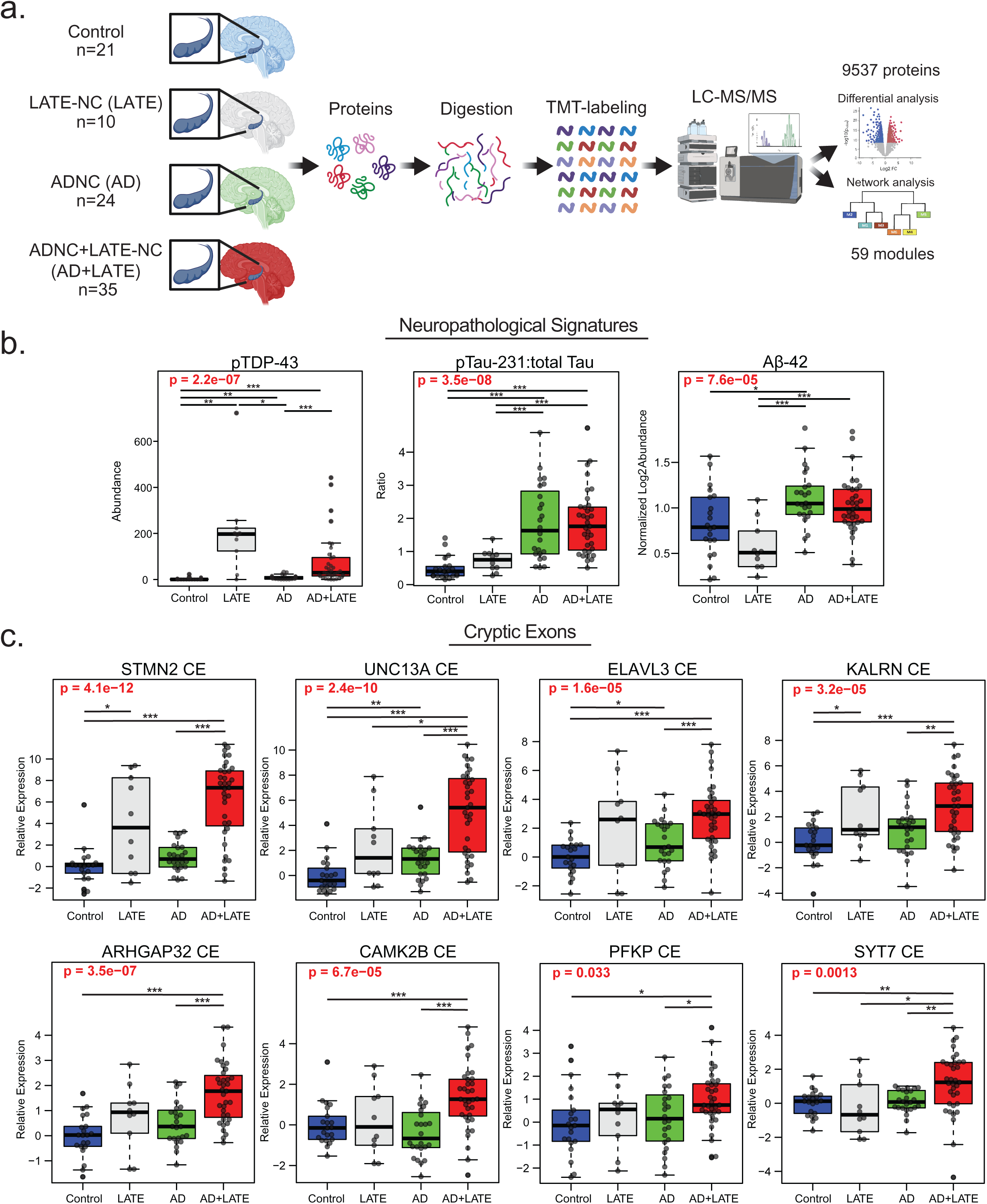
Proteomic workflow and biochemical/molecular measures of ADNC and LATE-NC biomarkers in postmortem hippocampus. A. Schematic of the experimental workflow for examining proteomic differences in postmortem hippocampal tissue from 90 individuals classified into four groups: controls (n = 21), ADNC without LATE-NC (n = 24), ADNC+LATE-NC (n = 35), and LATE-NC (n = 10). B. Boxplots showing immunoassay levels for pTDP-43, pTau-231 relative to total Tau, and TMT-MS abundances for Aβ42. The ratio of phosphorylated Tau (T231) to total Tau is presented as log₂ abundance, and Aβ levels were quantified using TMT-MS following quality control. C. Boxplots depicting log₂ abundance of cryptic exons (CEs) determined by qPCR for STMN2, UNC13A, ELAVL3, KALRN, CAMK2B, PFKP, SYT7, and ARHGAP32. Analyses of variance (ANOVA) results are highlighted in bold red when disease type significantly influenced protein/peptide abundance. Lines at the top of each plot indicate individual t-tests for direct comparisons, with significance denoted by asterisks (*p < 0.05, **p < 0.01, ***p < 0.001).

To first investigate the biochemical impact of AD and LATE pathology, we performed immunoassays and mass spectrometry to quantify key disease-associated proteins, including phosphorylated TDP-43 (pTDP-43), phosphorylated Tau at threonine 231 (pTau231) to total Tau ratio, and Aβ42 (measured by TMT-MS as described^60^) across control, AD, LATE, and AD+LATE subgroups (**Figure 1B**). pTDP-43 levels varied significantly across disease groups (p = 2.2e−07), with the highest levels observed in LATE, followed by AD+LATE consistent with the neuropathological classification (**Supplemental Table 1**). Although AD cases exhibited slightly higher relative pTDP-43 levels compared to controls, these levels were still significantly lower than those in LATE and AD+LATE. In contrast, AD and AD+LATE cases showed increased Tau phosphorylation levels compared to controls and LATE, though no significant differences were observed between AD and AD+LATE. Aβ42 abundance also varied across groups (p = 7.6e−05), with significantly higher levels in AD compared to controls and LATE, while AD+LATE showed comparable Aβ-42 levels to AD. Interestingly, LATE cases had lower Aβ42 levels, consistent with the expectation that Aβ plaque and tau neurofibrillary tangle (NFT) pathology are primarily associated with AD rather than LATE neuropathological changes^61^. These biochemical findings support the pathological assessment that pTDP-43 pathology is most prominent in cases with LATE and AD+LATE.

To evaluate the impact of AD and LATE pathology on cryptic exon (CE) inclusion that results from TDP-43 dysfunction, we quantified the relative expression of CE-containing transcripts across control, AD, LATE, and AD+LATE groups. CE inclusion was assessed in transcripts previously associated with TDP-43 dysfunction, including CEs in *STMN2*, *UNC13A*, *ELAVL3*, *KALRN*, *CAMK2B*, *PFKP*, *SYT7*, and *ARHGAP32*, all of which showed significant alterations across disease groups (**Figure 1C**). *STMN2* and *UNC13A* CE inclusion exhibited the most pronounced increases in the LATE and AD+LATE groups compared to controls and AD cases, consistent with the presence of TDP-43 pathology. Similarly, *ELAVL3* and *KALRN* CEs were significantly elevated in AD+LATE and LATE, with the strongest effects observed in AD+LATE (p = 1.6e−05 and p = 4.6e−10, respectively). Additional CEs in *CAMK2B*, *PFKP*, *SYT7*, and *ARHGAP32* also showed significant disease-related differences (ANOVA p < 0.05 for all comparisons), with higher levels of CE inclusion in AD+LATE cases. As expected, CE inclusion was present, but generally lower in AD cases compared to AD+LATE consistent with previous reports^62^. However, in some instances, particularly for *UNC13A* and *ELAVL3*, CE expression was significantly increased in AD compared to controls, which may be related to a modest increase in pTDP-43 levels by immunoassay in AD cases and could explain the higher levels of CE observed in AD+LATE. These findings highlight the complex relationship between TDP-43 and AD pathologies and their impact on TDP-43 mis-splicing and CE inclusion.

### Relationship Between Cryptic Exon Inclusion and Pathological Hallmarks of AD and LATE

To further examine the relationship between CE inclusion due to TDP-43 dysfunction and key pathological markers of AD and LATE, we performed a pairwise correlation analysis between CE-containing transcripts (*STMN2*, *UNC13A*, *ELAVL3*, *ARHGAP32*, *CAMK2B*, *KALRN*, *PFKP*, and *SYT7*) and disease-associated pathologic proteins, including pTDP-43 alone, pTau/Tau ratio, and Aβ42 (**Figure 2A**). We observed the strongest positive correlations between CE transcripts and pTDP-43, particularly for *STMN2*, and *UNC13A*, which exhibited the highest correlation coefficients (Cor ∼ 0.45 with p<0.001). These findings further reinforce the role of TDP-43 dysfunction in cryptic splicing events. In contrast, correlations between the measured CE transcripts and pTau/Tau ratio or Aβ42 levels were generally weaker. While some CE transcripts, such as *STMN2* and *UNC13A*, showed modest positive associations with pTau pathology (cor = 0.31 and 0.29; respectively), these relationships were generally weaker across all CEs measured (average Cor = 0.17) than those observed with pTDP-43 pathology (average Cor = 0.37). Aβ42 levels exhibited minimal to no correlation with CE expression (Average Cor = -0.01) indicating that CE inclusion is primarily linked to pTDP-43 pathology rather than AD-related amyloid accumulation. We also performed pairwise correlation analysis among CE-containing transcripts to assess their co-regulation across different genes (**Figure 2B**). Strong positive correlations were observed between several CEs, suggesting a shared regulatory mechanism, driven by TDP-43 loss of function. *STMN2* CE and *UNC13A* CE exhibited the highest correlation (Cor = 0.87), consistent with their established roles as downstream targets of TDP-43 splicing regulation. Similarly, CEs in *ELAVL3*, *CAMK2B*, *ARHGAP32*, and *KALRN* displayed moderate to strong correlations (Cor = 0.62–0.72) to each other, whereas CEs in *PFKP* and *SYT7* showed weaker correlations with other cryptic exons (Cor = 0.39–0.58), indicating, additional factors may influence their splicing regulation in disease. These findings suggest that CE inclusion is a tightly co-regulated process affecting multiple genes, with certain CE transcripts, such as *STMN2* and *UNC13A*, being more strongly associated. Importantly, while CE inclusion is, as expected, positively correlated with pTDP-43, RNA transcripts harboring CEs exhibit stronger correlations with each other than with pTDP-43, indicating that TDP-43 aggregation (gain of function) and cytoplasmic mis-localization (loss of function) are not always co-dependent.

**Figure 2.**
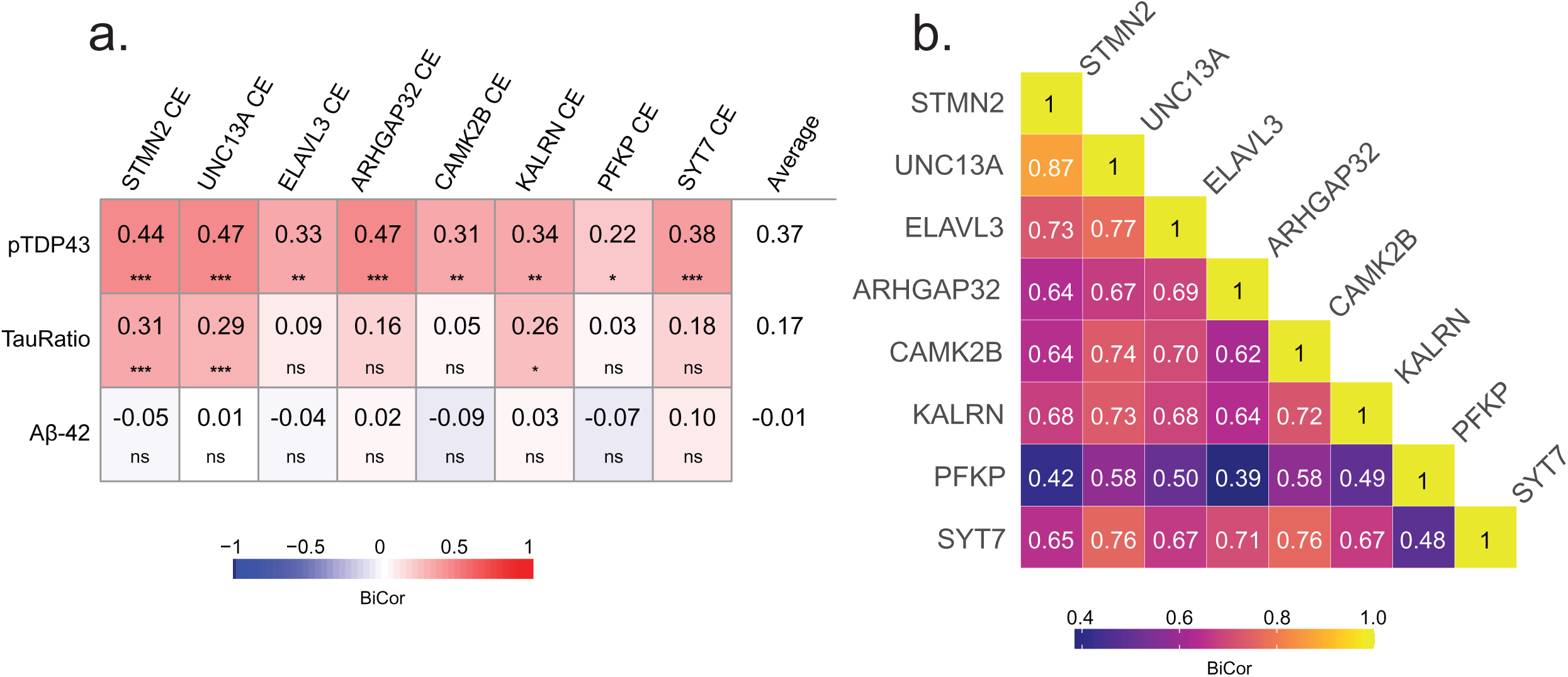
Cryptic exons are more strongly associated with each other than with pathological markers pTDP-43, pTau231 ratio, and Aβ42. A. CE levels of STMN2, UNC13A, PFKP, ARHGAP32, CAMK2B, SYT7, ELAVL3, and KALRN are correlated with pathological markers pTDP-43, pTau-231, and Aβ42. B. Correlations between individual CEs. Heatmaps display biweight midcorrelations (BiCor) values between each CE and relevant pathological markers, as well as pairwise correlations among CEs. Student correlation p-values for multiple biweight midcorrelations are indicated by asterisks (*p < 0.05, **p < 0.01, ***p < 0.001).

### Molecular subtyping based on CE burden across the spectrum of LATE and Alzheimer’s Disease

Since pTDP-43 levels and CE harboring transcripts are not as strongly correlated compared to the high correlation observed among CE transcripts themselves, we sought to unbiasedly cluster individual cases into low, intermediate, and high CE burden subtypes. This hierarchical clustering was based on the expression profiles of the eight CE-containing transcripts across all 90 hippocampal tissues (**Figure 3A**). Principal component analysis (PCA) confirmed the validity of this classification, showing clear separation between low and high CE burden groups, with an intermediate group occupying in between (**Supplemental Figure 2a**). Notably, individuals from each pathological subgroup (Control, LATE, AD, and AD+LATE) were represented across all CE burden subtypes (low, intermediate, and high), except for AD-only cases, which were absent from the high CE burden group (**Figure 3B**). The highest percentage of individuals in the high CE burden subtype were AD+LATE cases, whereas control and AD cases were predominantly classified in the low CE burden group, with some exceptions. For example, two control cases was deemed to have intermediate and high CE burden, whereas some LATE cases were determined to have low CE burden. This distribution of pathologically defined cases further highlights the specificity of the CE subtypes. As expected, the cumulative or aggregate CE burden score was significantly different across subtypes, showing strong statistical support (p = 2.7e−30) (**Figure 3C**). Additionally, Mini-Mental State Exam (MMSE) cognitive scores collected prior to death were significantly lower in individuals with high CE subtypes compared to those with low CE subtypes, indicating that a higher CE burden is associated with greater cognitive dysfunction (**Supplemental Figure 2b**).

**Figure 3.**
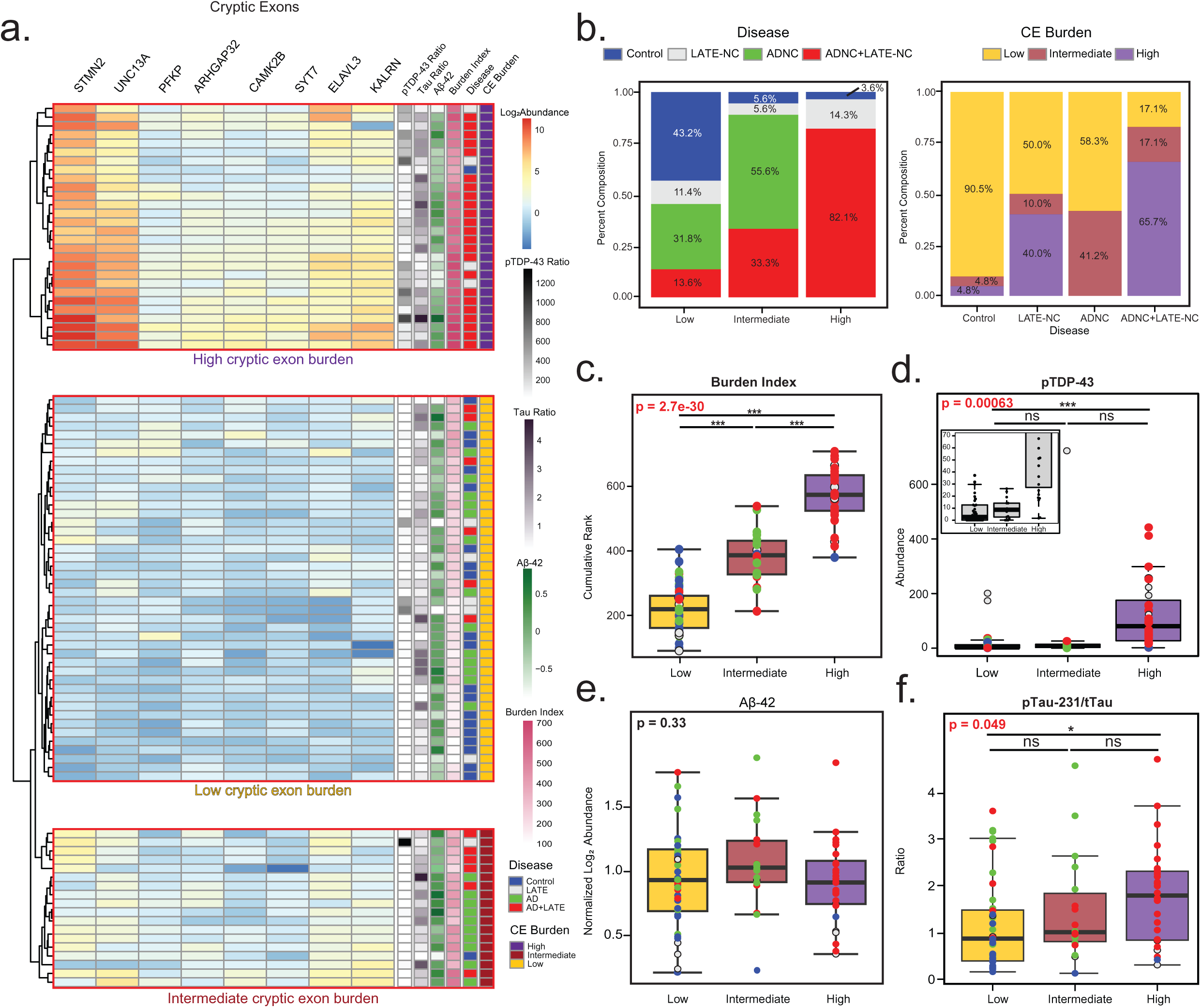
Hierarchical clustering of cryptic exon molecular profiles identifies distinct subtypes independent of ADNC pathological markers. A. Unbiased hierarchical clustering of log₂ abundance levels for eight common cryptic exons (STMN2, UNC13A, PFKP, ARHGAP32, CAMK2B, SYT7, ELAVL3, and KALRN) identified three CE subtypes: low, intermediate, and high. The heatmap displays relative CE abundance, with pTDP-43 (assay)/total TDP-43 (MS), pTau-231/total Tau, and Aβ42 log₂ abundance for each individual shown on the right. Additionally, cryptic exon burden, calculated as the cumulative rank of each individual’s CE abundance, is represented alongside pathological group classification. B. Percent composition of pathology subgroups (Controls, LATE-NC, ADNC, and ADNC+LATE-NC) within each CE subtype (high, intermediate, and low). Conversely, the distribution of CE subtypes across the pathological subgroups is also shown. C. Cumulative cryptic exon burden levels were stratified by high, intermediate, and low CE subtypes. D-F. pTDP-43, pTau-231, and Aβ42 levels were also analyzed by CE subtype. Analyses of variance (ANOVA) results are highlighted in bold red where cryptic burden significantly affected abundance. Lines at the top of each plot indicate individual t-tests for direct comparisons, with significance denoted by asterisks (*p < 0.05, **p < 0.01, ***p < 0.001).

Next, we examined how CE subtypes differed with key pathological markers (pTDP-43, pTau231/Tau ratio, and Aβ42). A strong association was observed between CE burden and TDP-43 pathology, with pTDP-43 significantly increased in individuals with high CE subtypes compared to those with low or intermediate burden (**Figure 3D**, p = 2.7e−30). However, the intermediate CE subtype, had lower levels of pTDP-43 compared to the high CE subtype. Although not statistically significant due to a single outlier, pTDP-43 levels in the intermediate CE subtype were comparable to those in the low CE subtype. This indicates that CE inclusion, driven by TDP-43 dysfunction, can occur even without significantly elevated biochemical levels of pTDP-43. Conversely, we also observed cases pathologically classified as LATE with high pTDP-43 levels, but that were classified to the low CE subtype. When assessing hallmark AD pathologies, Aβ42 levels did not differ significantly across CE subtypes (**Figure 3E**), further supporting that plaque pathology is not strongly linked to CE inclusion. Moreover, only a moderate increase in pTau231 was observed in cases classified as high CE subtype compared to the low CE subtype (p<0.05), likely due to the high proportion of AD+LATE cases in this group (**Figure 3F**). However, given the number of AD cases in the low CE subtype, hallmark AD pathologies (e.g. plaques and tangles) are not a primary driver of subtype classification. In summary, CE subtypes were strongly linked to TDP-43 pathology, with higher pTDP-43 levels in the high CE subtype, though CE inclusion could still occur in intermediate and even some high CE subtypes without elevated pTDP-43. In contrast, Aβ42 and pTau231 showed little to moderate increases with CE burden, suggesting that AD pathology do not strongly influence CE subtype classification.

### Global proteomic differences across cryptic exon subtypes

To investigate proteomic changes associated with CE burden, we performed differential abundance analysis comparing high, intermediate, and low CE subtypes. Across these comparisons, we identified widespread proteomic alterations, with numerous proteins showing significant increases or decreases in abundance depending on subtype (**Figure 4A-C and Supplemental Table 4**). When comparing high versus low CE subtypes, we identified 750 proteins with decreased abundance and 584 with increased abundance, highlighting extensive proteomic shifts in the high CE burden group. The most significantly increased Gene Ontology (GO) terms were associated with extracellular matrix (ECM) organization and integrin signaling, while the most decreased terms predominantly mapped to endosomal processes, endoplasmic reticulum (ER) function, and microtubule biology (**Supplemental Table 9**). In contrast, the intermediate versus low CE burden comparison revealed fewer differentially abundant proteins, with 235 decreased and 210 increased, suggesting that intermediate CE burden represents a transitional state with moderate proteomic alterations. Finally, the comparison between high and intermediate CE burden groups showed the fewest changes, with only 33 proteins decreased and 57 increased, indicating that these subtypes at the proteome level are relatively similar to each other. Consistently, we observed a positive correlation (Cor = 0.63, p = 1.1e-174) between proteomic changes in the intermediate and high CE burden subtypes relative to the low CE subtype, reinforcing the notion that high and intermediate CE subtypes share similar proteome alterations (**Figure 4D**). These differences across CE subtypes contrast with proteomic changes observed among pathological subtypes (Control, LATE, AD, and AD+LATE), where over 1,000 proteins exhibited increased or decreased abundance compared to controls.

**Figure 4.**
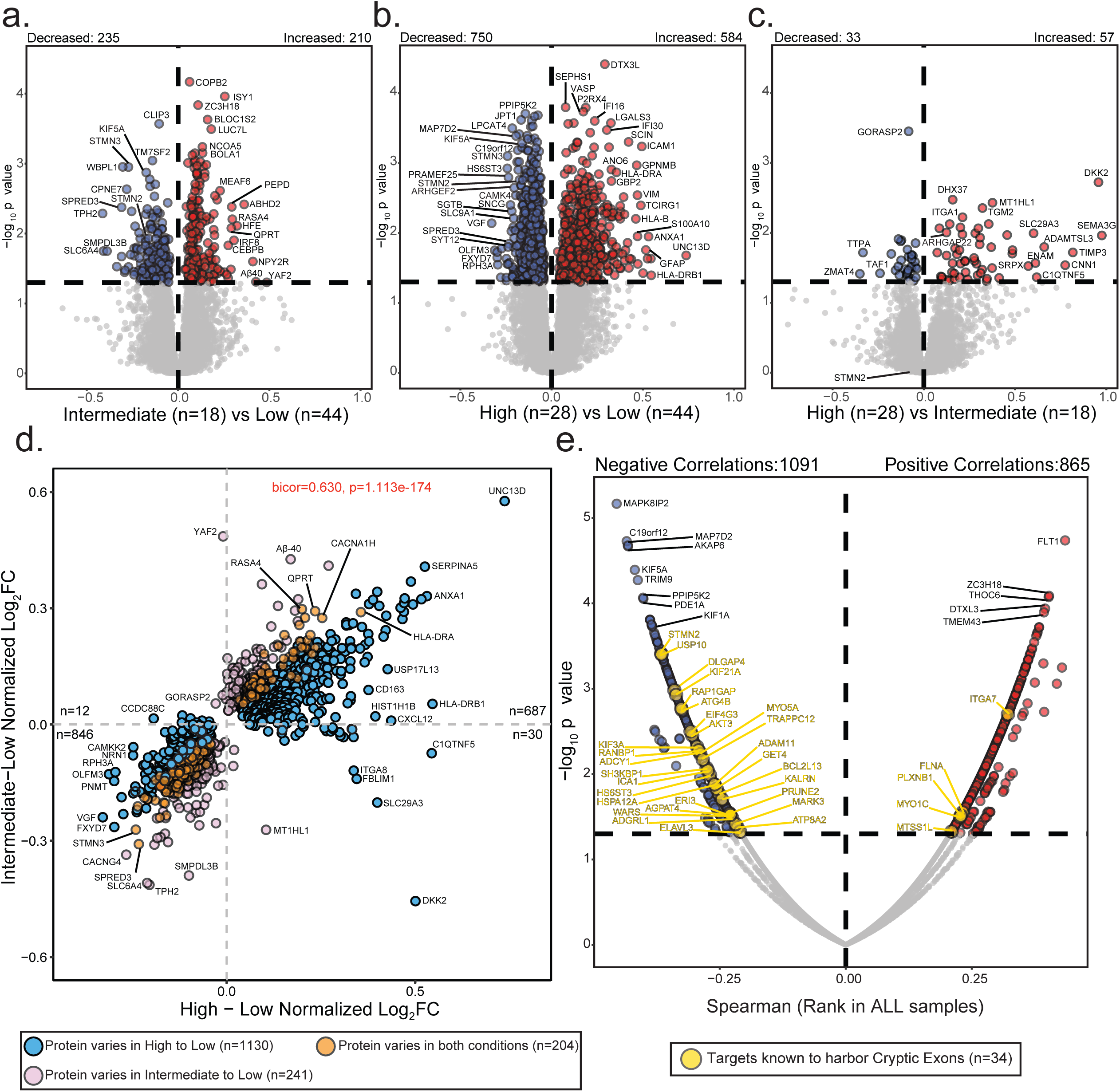
Comparative proteomic analyses across cryptic exon subtypes. A-C. Volcano plots depicting differential abundance profiles of hippocampal proteomes between cryptic exon (CE) subtypes: intermediate (n=18) vs. low (n=44), high (n=28) vs. low, and high vs. intermediate. Proteins with increased or decreased abundance are highlighted at the top of each panel. Log₂ fold change (x-axis) represents the magnitude of change, while one-way ANOVA with Benjamini-Hochberg corrected -log₁₀ p-values (y-axis) indicates statistical significance. Proteins significantly increased in abundance (p < 0.05) are shown in red, significantly decreased proteins in blue, and non-significant proteins in gray. D. Scatter plots comparing protein abundance changes between intermediate vs. low CE burden and high vs. low CE burden across 9,537 proteins. Blue points indicate proteins significantly different in the high vs. low comparison but not in intermediate vs. low. Pink points are significantly different in intermediate vs. low but not in high vs. low. Orange points represent proteins significantly altered in both comparisons. E. Spearman’s rank correlation between cryptic exon burden and protein abundance. The top correlated proteins are labeled. Gold points represent proteins whose corresponding transcripts are predicted to harbor cryptic exons.

These widespread changes in the proteome were largely driven by pathological burden (β-amyloid, tau, and TDP-43) across the clinicopathological subgroups resulting in well established “cellular phase” changes in the bulk proteome typically associated with AD^44, 45, 63, 64^ **Supplemental Figure 3 and Supplemental Table 4**). For example, plaque-associated proteins such as MDK, ICAM1, and NTN1, as well as the APOE4 isoform, were significantly increased in AD and AD+LATE cases compared to controls, consistent with the higher plaque load and genetic risk of AD in these groups. Similarly, markers of reactive astrocytosis, including GFAP, VIM, and CD44, were elevated in LATE, AD, and AD+LATE cases, reflecting increased glial activation in these pathological conditions. Conversely, proteins associated with synaptic loss, such as VGF and NRN1, were decreased in LATE, AD and AD+LATE pathological subgroups, further supporting the link between neurodegeneration and reduced synaptic integrity typically seen in AD.

### Proteomic-wide association to cryptic exon burden in the hippocampus

To further identify specific proteins impacted by CE burden in the frontal cortex, we performed a correlation analysis across 9,537 pairwise protein comparisons, using cumulative CE burden scores from 90 unique donors (**Figure 4E**). Since CEs are non-conserved intronic sequences, they often introduce splicing errors leading to frameshifts, pre-mature stop codons, or alternative polyadenylation^26, 65^. These errors can reduce transcript expression through NMD or other RNA degradation pathways, often resulting in a corresponding decline in their encoded protein levels. For example, CE inclusion in *STMN2* leads to the loss of *STMN2* RNA and protein in ALS and FTLD-TDP tissues^27, 28, 31^. Based on these findings we hypothesized that protein levels of these CE transcripts would be reduced with increasing CE burden in individual hippocampal cases. To test this, we used RNA-seq evidence from two studies to generate a comprehensive list of 183 transcripts that harbor CEs due to TDP-43 dysfunction^30, 56^. We then examined the relationship between the encoded protein levels of these CE transcripts and cumulative CE burden in the hippocampus. Importantly, we observed a bias in protein levels, with 29 proteins encoded by CE harboring transcripts showing a negative correlation or reduction in cases with high CE burden, compared to only five proteins from CE transcripts that were positively correlated (**Figure 4E**). Proteins significantly negatively associated with CE burden included STMN2, ELAVL3, and KALRN, all of which had high levels of CEs within their transcript in these exact hippocampus tissues (**Supplemental Figure 4**). We also observed a significant reduction in kinesin proteins, including KIF3A and KIF21A, both of which have been identified to harbor CEs via RNA-seq^30, 56^ (**Supplemental Figure 5**). Interestingly, while *KIF3A* and *KIF21A* transcripts are known CE targets, protein levels of KIF1A and KIF5A showed an even greater reduction in cases with higher CE burden. Notably, mutations in *KIF1A* and *KIF5A* are known to cause rare familial forms of ALS ^66, 67^, suggesting a shared mechanism involving TDP-43 loss of function and kinesin dysregulation. There were five proteins from putative CE transcripts that increased with higher CE burden, including ITGAM and PLXNB1, which are predominantly expressed in microglia and astrocytes, respectively. This increase may result from an increased glial number or altered activity states due to neuroinflammation. In summary, we found that higher CE burden in the hippocampus is associated with a widespread reduction in protein levels encoded by CE-containing transcripts, particularly neuronal proteins such as STMN2, ELAVL3, KALRN, KIF3A, and KIF21A, supporting a model in which TDP-43 dysfunction leads to mis-splicing, RNA degradation, and reduced protein expression.

### Network analysis reveals shared and divergent modules associated with pathological and CE subtypes

To investigate system-level changes in the hippocampal proteome associated with CE burden and clinicopathological phenotypes, we performed Weighted Gene Co-expression Network Analysis (WGCNA)^57^ to identify co-expressed protein modules or communities of proteins correlated with CE burden, cell types, and neuropathological traits^68, 69^. WGCNA revealed distinct protein modules, ranging from the largest module (M1) with 482 proteins to the smallest (M59) with 24 proteins, leaving only 355 of 9,537 proteins unassigned. Several modules exhibited high interrelatedness, suggesting shared abundance patterns and biological functions. As expected, we identified modules associated with brain cell types, including a cluster (M3, M1, M12, M2, and M11) enriched in astrocyte and microglia markers, which are involved in extracellular matrix (ECM) organization and antigen binding. In contrast, oligodendrocyte and endothelial markers were predominantly expressed in M4 and M24, respectively, with roles in myelination and basement membrane biology. Neuronal modules were more broadly distributed across the network, including M5, M15, and M30, which were associated with synaptic transmission and GPCR signaling, as well as M9 and M10, which were involved in synaptic and dendritic biology.

Correlation analyses between protein modules and neuropathological traits revealed that pTDP-43 and pTau/total tau ratio exhibited similar correlation trends, aligning primarily with AD and AD+LATE pathological grouping (**Figure 5**). Similarly, correlations between modules demonstrated clustering of co-expression modules into disease relevant groups (**Figure 5A**). Specifically, we observed increased astrocyte and microglia module levels and a decrease in synaptic module levels in these subtypes. However, these module changes differed from those associated with Aβ42, likely due to the lower Aβ42 levels in LATE cases compared to controls (**Figure 1**). Controls in this study showed evidence of diffuse plaques^33^. However, LATE cases showed increased glial module expression and decreased neuronal module levels despite low pTau and plaque burden, suggesting that pTDP-43 pathology and/or CE inclusion alone may drive the cellular changes typically seen in AD and related dementias (**Figure 5B**).

**Figure 5.**
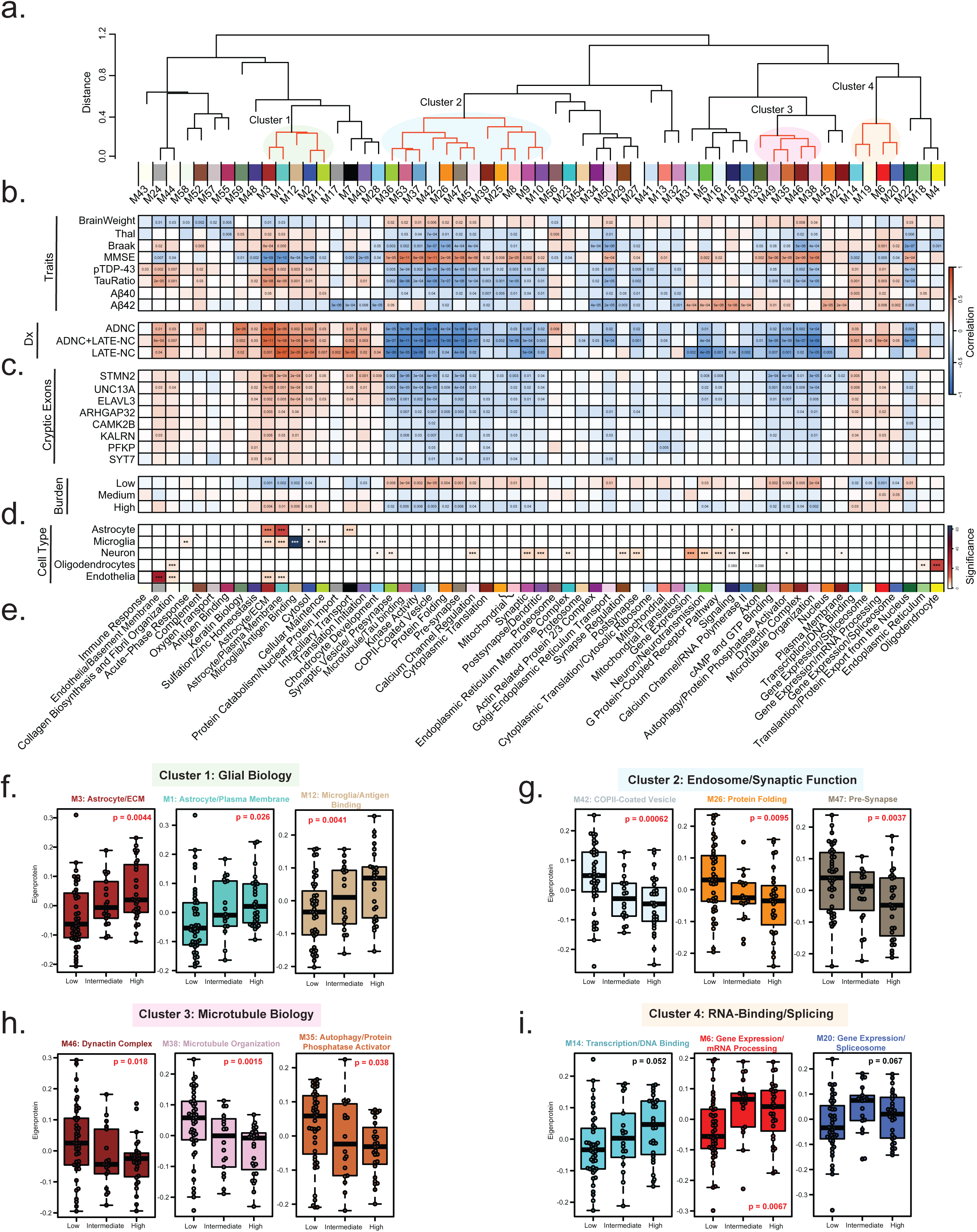
Network analysis of protein modules and their associations with pathology, cryptic exon subtypes, and cell-type enrichment. A. Cluster dendrogram illustrating the similarity of WGCNA network modules based on correlations between eigenproteins (first principal component of each module). Clusters of related modules are indicated by transparent colored circles (Clusters 1, 2, 3, and 4). B. Correlations between pathological, molecular, and clinical traits, including disease markers, and individual protein modules were assessed using Biweight midcorrelation (BiCor) analysis. Pathological diagnosis (ADNC, ADNC+LATE-NC, and LATE-NC) was evaluated through BiCor, comparing each disease subgroup separately versus control with module eigenproteins. C. Associations between eight cryptic exons and protein modules were evaluated using BiCor analysis of module eigenproteins. Cryptic exon burden subtypes (high, intermediate, and low) were analyzed similarly, assessing each with module eigenproteins. Significance levels from BiCor analysis are indicated by asterisks (*p < 0.05, **p < 0.01, ***p < 0.001). D. Cell-type enrichment was determined by comparing module proteins with reference lists of proteins enriched in astrocytes, microglia, neurons, oligodendrocytes, and endothelial cells (see Methods). Significance levels, determined by one-tailed Fisher’s exact test, are indicated by asterisks (*p < 0.05, **p < 0.01, ***p < 0.001). E. Top gene ontology (GO) terms were selected from significant GO annotations (**Supplemental Table 9**). F–I. Module eigenprotein levels are shown by subtype for three representative modules from each functional cluster (i.e., glial biology, endosome, microtubule biology, and RNA-binding proteome, respectively). P-values from one-way ANOVA are displayed, with significant values (α = 0.05) highlighted in bold red.

When compared to clinicopathological subtypes (control, LATE, AD, and AD+LATE), molecular CE subtypes exhibited both shared and distinct module differences. Examining the relationship between CE subtype, individual CEs, we found consistent associations across modules, highlighting those particularly linked to CE inclusion. In total, 34 modules were negatively correlated with CE burden, including the individuals CE transcripts (**Figure 5C**). The dendrogram revealed four main clusters of modules that showed either positive or negative correlations with CE burden (**Figure 5A**). Modules in Cluster 1 (M3, M1, M12, M2, and M11), including astrocytic and microglial markers (**Figure 5D**), were elevated, suggesting that a higher CE burden is associated with increased neuroinflammation. Additionally, Cluster 4 (M14, M19, M6 and M20) contained modules positively correlated with CE burden, as highlighted in the dendrogram. These modules included RNA-binding proteins, transcription factors, and splicing regulators (**Figure 5E**), notably including M6, which contained TDP-43 itself. This is particularly interesting, as TDP-43 mis-localization and loss of expression is thought to drive CE signatures. However, the observed increase in TDP-43 and other RNA-binding proteins in M6 with higher CE burden may reflect aggregation processes or a compensatory response to the pathological consequences of TDP-43 dysfunction.

Modules that decreased with higher CE burden were primarily grouped into Cluster 2 and Cluster 3, with some exceptions (**Figure 5F-I**). Cluster 2 (M36-M10), the largest cluster (n=12 modules), contained two major sub-clusters: 2A (M36-M5) and 2B (M39-M10). Each sub-cluster was associated with distinct biological processes, including vesicular transport, presynaptic function, calcium channel signaling, and protein folding. These modules were predominantly linked to neuronal cell types, consistent with CE burden impacting synaptic function. Similarly, modules in Cluster 3 (M33-M38) proportionally decreased with increasing CE burden across subtypes. These modules were functionally associated with cytoskeleton function including GTP binding, autophagy, dynein activity, and microtubule function. Although not part of a specific cluster, M5 (Synaptic Transmission), the largest neuron-enriched module, was also proportionally reduced across CE subtypes with increasing CE burden. Collectively these data support the hypothesis that CE inclusion impacts synaptic structure and function (i.e., vesicle transport, and cytoskeleton function).

### Integrated proteomics prioritizes CE targets and pathways in both hippocampal tissues and human neurons affected by TDP-43 insufficiency

To identify causal changes resulting from TDP-43 loss of function, we performed proteomics on CRISPRi TDP-43 depleted and control human iPSC derived neurons (iNeurons). In total, we identified 7,389 proteins, with 876 significantly increased and 610 significantly decreased in the TDP-43 KD iNeurons compared to control (**Figure 6A-B**). These TDP-43 deficient iNeurons were previously shown to harbor significant number of CE transcripts across a broad range of genes^32^. As expected, TDP-43 protein levels were significantly reduced (∼10-fold change, p value=3.4e-6) in the TDP-43 depleted iNeurons. Notably, we observed reduced levels of 91 percent of proteins derived from putative CE transcripts, while only 9 percent of proteins derived from putative CE transcripts showed an increase in abundance (highlighted in yellow) consistent with loss of expression due to pre-mature truncation, polyadenylation and/or NMD mechanisms. This included STMN2, KALRN, and ELAVL3, which were also reduced in high CE subtypes in the hippocampus. These data support previous findings showing that CEs within transcripts resulting from TDP-43 knockdown lead to reduced protein levels^32^. Using integrated transcriptomics and proteomics (i.e., proteogenomics) of TDP-43-depleted iNeurons, we previously identified peptides originating from *de novo* proteins encoded by cryptic exons^32^. Here, leveraging data independent acquisition-mass spectrometry (DIA-MS) coupled with the enhanced sensitivity of the Orbitrap Astral mass spectrometer we aimed to confirm these findings and identify novel mis-spliced CE transcripts that translate into *de novo* proteins. We leveraged custom databases containing *in silico* translated protein sequences of CEs^32^, incorporated into a standard human proteome reference, and applied an enhanced 5-minute liquid-chromatographic (LC) elution gradient to increase peptide signal intensities. We identified cryptic peptides from HDGFL2, RSF1, SYT7 and MYO18A, validating previous results^32^, and discovered a new in-frame cryptic peptide in DNM1 (**Supplemental Figure 6; Supplemental Table 10-11**). These identified peptides were predominantly found in TDP-43 KD iNeurons and, even when detected in control iNeurons, were several orders of magnitude lower in intensity (**Supplemental Figure 6**; **Supplemental Table 11**). With the exception of MYO18A, the only cryptic peptide uniquely detected using the 30-minute LC gradient, all other cryptic peptides showed a greater than 8-fold increase in TDP-43 KD iNeurons compared to controls. In control samples, the majority of cryptic peptide intensities were imputed due to missing values, indicating low or undetectable levels under normal conditions. Overall, our proteomic analysis of TDP-43 loss of function in human neurons revealed widespread reductions in protein encoded from CE transcripts, confirming their instability, while enhanced mass spectrometry identified both known and new CE-derived peptides, further linking TDP-43 insufficiency to altered changes in the proteome.

**Figure 6.**
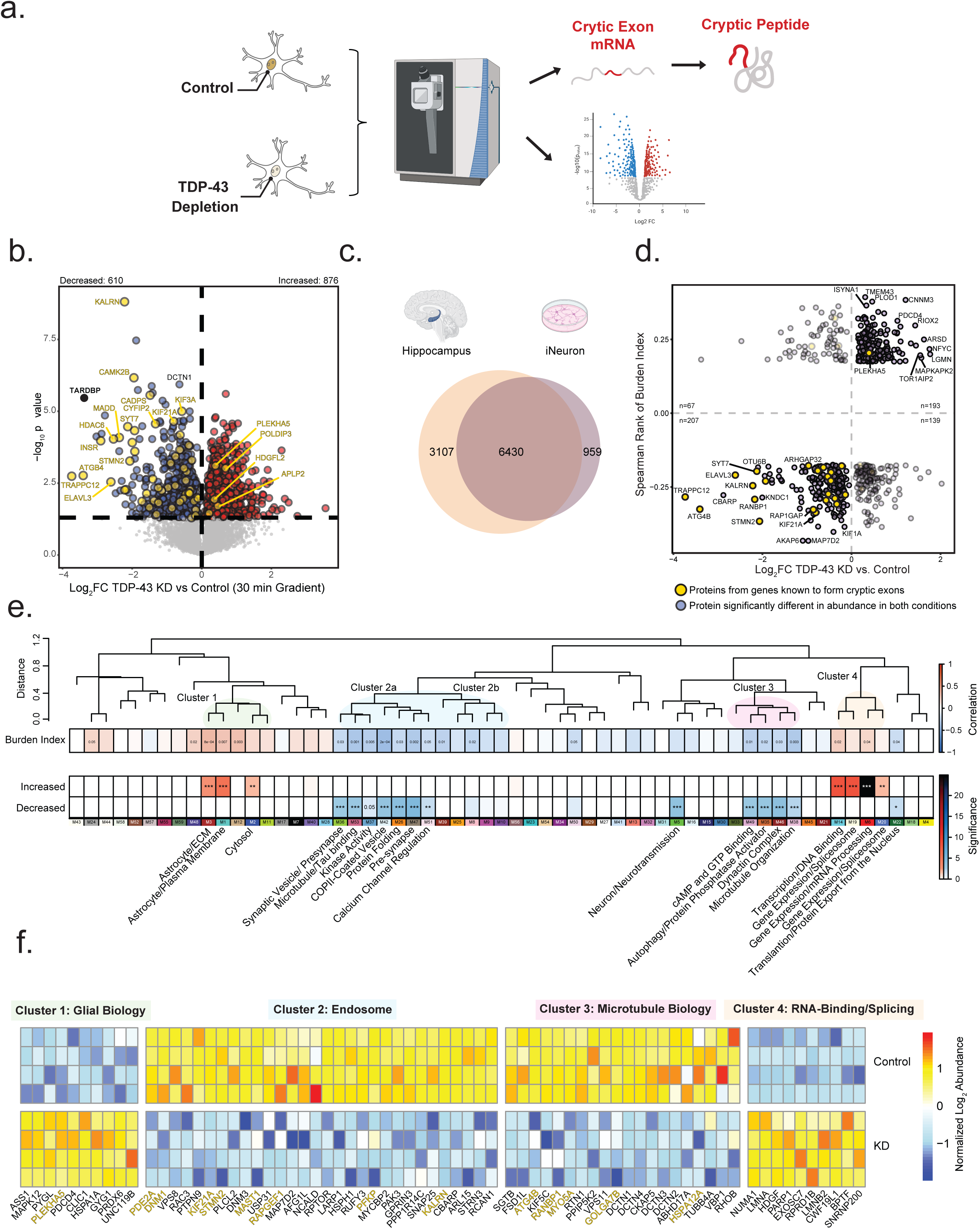
Overlap between human hippocampal network modules and differentially abundant proteins in TDP-43 knockdown iNeurons. A. Schematic of the experimental workflow used to examine proteomic differences between TDP-43 knockdown (KD) and control iNeurons. B. Volcano plots depicting differential protein abundance between TDP-43 KD and control iNeurons using a 30-minute liquid chromatography (LC) gradient. A total of 7,389 proteins were identified, with 876 increased and 610 decreased in abundance. Log₂ fold change (x-axis) represents the magnitude of change, while one-way ANOVA with Benjamini-Hochberg corrected -log₁₀ p-values (y-axis) indicates statistical significance. Proteins significantly (p < 0.05) increased in abundance are outlined in red, significantly decreased proteins in blue, and non-significant proteins in gray. C. Venn diagrams illustrating the overlap between the neuronal proteome (n=7,389) and postmortem hippocampal proteome (n=9,537). D. Scatter plot showing proteins overlapping at α = 0.1 between neurons (TDP-43 KD vs control) and Spearman’s correlation of cumulative cryptic exon (CE) burden score. Proteins encoded by transcripts predicted to harbor CEs are highlighted in gold. E. CE burden score was correlated with eigenprotein values using BiCor analysis. Additionally, proteins concordant in direction of change between TDP-43 knockdown vs. control neurons and Spearman’s correlation of the CE Burden score were compared to module membership in the hippocampal proteome. Significance levels, determined by one-tailed Fisher’s exact test, are denoted by asterisks (*p < 0.05, **p < 0.01, ***p < 0.001). A dendrogram of module relatedness is shown, with related modules highlighted by transparent circles. Gene ontology (GO) terms for relevant modules are listed below. F. Heatmap of differentially abundant proteins in iNeuron cultures. Five representative proteins from each module enriched in concordant proteins between postmortem hippocampus and iNeurons are shown. Heatmap is divided by condition (control vs. TDP-43 KD) and biological cluster. Proteins from cryptic exon (CE)–containing genes are highlighted in gold.

### Proteomic Signatures of TDP-43 Deficiency Reveal Concordant Changes in CE Burden and Brain Network Modules

To validate protein products of CE transcripts with reduced expression due to TDP-43 dysfunction in human tissues, we compared hippocampal proteins that were positively and negatively correlated with the CE burden index score to proteins increased or decreased in the TDP-43 KD neuronal proteome, analyzing 6,430 overlapping proteins (**Figure 6B-C**). We found that most proteins (nominally significant in both datasets FDR p value <0.1) showed concordant changes, with 400 proteins displaying a similar direction of effect (**Figure 6D**). Proteins consistently reduced with higher CE burden in tissue and TDP-43 KD neurons included STMN2, ELAVL3, KALRN, SYT7, and ARHGAP32, all of which had measurable CEs in the hippocampus. To better understand the biological connection between proteins reduced due to TDP-43 loss of function in human neurons and brain network modules, we performed a comprehensive integrated analysis. Specifically, we used Fisher’s exact test (FET) to determine which brain network modules significantly overlapped with proteins that were concordantly increased or decreased due to TDP-43 deficiency. We further categorized these modules into previously described clusters based on their expression trends across CE subtypes (**Figure 6E**). Proteins that were concordantly decreased in TDP-43 KD cell lines and that were reduced with high CE burden (blue) overlapped with 12 brain modules. Interestingly, these decreased proteins mapped to a subset of Cluster 2 modules (M36-M31) with biology predominantly involved in endosomal and synaptic functions. This is further highlighted by the individual protein overlap between TDP-43 KD proteins and M42 (COPII-coated vesicles), M26 (protein folding), and M47 (pre-synaptic proteins). PFKP (M26) and TRAPPC12 (M42) were among the most significantly reduced proteins in human neurons following TDP-43 KD, both of which have been identified at the RNA level to harbor CEs due to loss of TDP-43 function^30, 56^. Similarly, modules in Cluster 3 (M49-M3), associated with microtubule biology, showed significant overlap with proteins that were reduced in TDP-43 KD neurons. Two modules outside these clusters, M5 (synaptic transmission) and M18 (endoplasmic reticulum), also showed significant overlap. Both modules were negatively associated with CE burden and contained proteins that were significantly reduced following TDP-43 KD.

Proteins that were concordantly increased in TDP-43 KD neurons and in tissues with high CE burden (red) overlapped with 7 brain modules (**Fig. 5E and Fig. 6E-F**). This includes three modules in Cluster 1 (M3, M1, and M2), which are predominantly enriched with glial markers in the brain. Given that TDP-43 knockdown was performed in human neurons, this suggests that neurons may be at least partly responsible for the increased levels of these proteins observed in intermediate and high CE subtypes in the hippocampus. Additionally, all four RNA-binding/splicing modules in Cluster 4 (M14, M19, M6, and M20) were significantly enriched with proteins that increased following TDP-43 KD. Remarkably, in M6 (mRNA processing), all proteins, except for TDP-43 itself, were significantly increased following TDP-43 KD in the neurons, and this module was elevated in both intermediate and high CE subtypes in the hippocampus. The broad increase in RNA-binding proteins observed in human tissues with high CE burden and following TDP-43 KD suggests that TDP-43 deficiency is associated with a compensatory upregulation of the RNA-binding proteome.

## Discussion

Using an integrated proteomics approach on postmortem hippocampal tissues across a spectrum of LATE-NC and ADNC, we identified widespread proteomic changes associated with TDP-43 dysfunction. We found that eight TDP-43-regulated CEs correlate more strongly with each other than with detected pTDP-43 pathology. This suggests that CE inclusion is not always linked robustly with TDP-43 aggregation, and/or the “window” of detectable pTDP-43 pathology does not represent fully the TDP-43 dysregulation in the brain. Conversely, we also identified cases with high pTDP-43 levels but low CE burden, further supporting the dissociation between TDP-43 aggregation and cryptic exon inclusion. Therefore, to better capture proteomic changes linked to TDP-43 dysfunction, we applied a molecular subtyping approach, classifying cases into low, intermediate, and high CE burden subtypes, independent of pathological and clinical diagnosis of ADNC and LATE-NC. Higher CE burden was associated with cognitive decline and widespread proteomic alterations. Proteomic analysis revealed significant reductions in proteins encoded by CE-containing transcripts, including STMN2, ELAVL3, and KALRN, as well as kinesin proteins such as KIF5A and KIF1A, which are causal genes in ALS. Network analysis further showed that higher CE burden was associated with increased glial/neuroinflammatory modules and decreased modules related to endosomal/vesicle, autophagy, protein folding and synaptic pathways. Finally, proteomic profiling of human iPSC derived neurons following significant TDP-43 knockdown validated that CE inclusion leads to reduced protein levels and confirmed the translation of specific cryptic exons at the protein level. This study further supports CE burden as a molecular hallmark of TDP-43 loss of function with potential implications for protein biomarkers linked to TDP-43 dysfunction.

Our initial intent was to define proteomic changes linked to the clinical-pathological subtypes of LATE-NC, ADNC, and ADNC+LATE-NC, which are provided. However, we were intrigued by the stronger correlations among CEs compared to their correlation with biochemical levels of detected pTDP-43. While *STMN2* and *UNC13A* CEs were significantly correlated with pTDP-43 pathology as expected^62, 70^, this association was notably weaker than the correlation among CEs themselves. The correlation between pTDP-43 pathology and CE inclusion (BiCor ∼0.4) is consistent with other studies^70^ and may reflect the presence of CE burden in some ADNC cases that lack high levels of pTDP-43 pathology as reported previously^66 70, 71^. Conversely, we observed the opposite pattern in certain LATE-NC cases, which exhibited very high pTDP-43 levels. This reinforced the observation that TDP-43 pathology and CE burden are not always co-dependent and prompted our unbiased molecular subtyping based on the relative expression of eight CEs across tissues. These CE subtypes, which were independent of amyloid and tau pathology, exhibited distinct pTDP-43 profiles, with only the high burden CE subtype showing elevated pTDP-43 levels. The enrichment of ADNC cases within the intermediate, but not the high CE subtype, supports that TDP-43 loss of function may precede overt TDP-43 aggregation^19 18^. Therefore, a reasonable hypothesis is that individuals with ADNC in the intermediate CE subtype would have progressed to ADNC+LATE-NC had they lived longer, accumulating a greater burden of CE inclusion along with more evident cytoplasmic pTDP-43 aggregation. Collectively, these data from human postmortem tissues highlight the heterogeneity of TDP-43 pathophysiology and support biomarker studies to detect CE inclusion prior to symptom onset.

CE inclusion can lead to reduced transcript expression through mechanisms such as pre-mature truncation, alternative polyadenylation, or NMD, ultimately resulting in protein loss of function^26^. However, some CEs can be translated, giving rise to *de novo* cryptic peptides, which may exhibit a gain-of-function effect^19, 32, 72^. While we validated known and new cryptic peptides via mass spectrometry for HDGFL2, RSF1, SYT7, DNM1, and MYO18A in human iPSC derived neurons following signification TDP-43 knockdown (∼10 fold)^32^, detecting translated CEs at the peptide level using discovery-based mass spectrometry approaches in complex matrices like brain tissue remains particularly challenging, especially when not all cells exhibit loss of TDP-43 localization or overt pathology. In contrast, targeted assays for HDGFL2 cryptic neo-epitopes show robust increase in cases with TDP-43 pathology in both brain, plasma and CSF, although for many samples there was observable signal in controls^19, 32, 72^. To this end, after re-searching the TMT-MS raw data from the hippocampus against a custom database with translated CE sequences, the only cryptic peptide that passed FDR thresholds was MYO18A, which did not differ significantly across CE subtypes (p=0.57). While new tools and more specific antibodies are being developed to better detect translated CEs, our findings suggest that, for certain gene products, the corresponding reduction in full-length protein levels, such as STMN2, KALRN, ELAVL3, and kinesin family proteins, could serve as surrogate markers of TDP-43 dysfunction, given their consistently reduced expression in affected tissues. Future studies examining the relationship between these protein reductions, TDP-43 loss of function, and CE inclusion in paired CSF, plasma, and tissue samples from the same individuals will be needed to validate this hypothesis.

One strength of our systems level proteomic network analysis is its ability to identify both distinct pathways and shared biological processes that are most positively or negatively impacted by CE burden in the brain. STMN2 is essential for normal axonal outgrowth and regeneration and when a CE is included in *STMN2*, it results in a truncated form of the protein, leading to neurodegeneration^27, 31^. In the hippocampal proteomic network, STMN2 is a hub protein in module M53, whose members are functionally involved in microtubule dynamics and tau binding. This module was significantly decreased in cases with higher CE burden and included proteins that were also reduced following TDP-43 knockdown in iNeurons. This suggests that the majority of the module members linked to STMN2 are collectively decreased with higher levels of CE burden corresponding to TDP-43 dysfunction. Similarly, KALRN and DLGAP4 are also CE targets, both involved in synaptic function and post-synaptic density and map to modules M51 and M5, that are decreased with higher CE burden in brain and enriched with proteins that are significantly reduced following TDP-43 knockdown. Therefore, a major consequence of TDP-43 loss of function is a cascade of mis-splicing events, which contribute to synaptic dysfunction and ultimately cognitive decline. Supporting this, we found that individuals with high CE burden had significantly lower cognitive scores before death compared to those with low CE burden in the hippocampus, despite no significant differences in amyloid or tau pathology. These results suggest that co-occurring pTDP-43 pathology and CE inclusion directly impact cognitive performance. This aligns with previous studies showing that individuals with eventual autopsy-proven LATE-NC experience memory decline, even in the absence of AD pathology, indicating that TDP-43 pathology independently contributes to cognitive impairment^73^. We also observe a concomitant increase in network modules associated with microglial function and neuroinflammation in individuals with high CE burden. Many of these proteins are also significantly increased in neurons following TDP-43 knockdown, suggesting their expression may be more broadly shared across brain cell types. Future studies examining the cell-type–specific context of TDP-43 loss of function beyond neurons will be informative in assessing the specificity of CE inclusion across distinct genes and cellular populations.

Finally, brain modules enriched in RNA-binding proteins, including TDP-43 in module M6, were elevated in tissues with high CE burden. This is particularly notable given that TDP-43 aggregates and becomes phosphorylated in disease, and the subtypes with the highest pTDP-43 pathology also exhibited the greatest CE inclusion. We propose two possible, and potentially synergistic, explanations for the increased RNA-binding and splicing proteins observed in cluster 4 of human tissue. First, the increase may be driven by TDP-43 gain-of-function co-aggregation effects involving similar RNA-binding proteins, resulting from its insolubility. Alternatively, TDP-43 dysfunction resulting from cytoplasmic mislocalization may trigger a compensatory increase in the RNA-binding proteome, independent of aggregation, an effect supported by the TDP-43 knockdown results. Together, findings from both tissue and cell models underscore the broad impact of TDP-43 loss on the RNA-binding proteome, ultimately altering alternative splicing and downstream protein changes.

While this study provides important insights into the proteomic consequences of TDP-43 dysfunction and CE burden in both human hippocampal tissue and TDP-43-depleted iPSC-derived neurons, several limitations should be considered. First, the sample size for certain groups, particularly LATE-only cases, was relatively small, which may limit statistical power and the generalizability. However, this limitation is mitigated by our approach of subtyping all cases based on CE burden rather than relying solely on neuropathological classification. Second, while CE burden was strongly associated with pTDP-43 pathology, we cannot fully disentangle the contributions of TDP-43 aggregation (gain-of-function) versus nuclear depletion (loss-of-function) from this cross-sectional postmortem data alone. Additionally, CE expression and protein-level effects may be influenced by cell type composition, RNA stability, or regional vulnerability, which are not fully captured by bulk tissue proteomics employed here. Third, although the use of TDP-43 knockdown in iPSC derived neurons provides mechanistic support for causal relationships, these *in vitro* models lack the cellular diversity of the brain microenvironment, particularly glial interactions, which may be critical in shaping disease-related phenotypes. Finally, while we validated known and identified novel cryptic peptides using enhanced DIA-MS in iNeurons following TDP-43 knockdown, the detection sensitivity for low-abundance or potentially unstable CE peptides from these proteins in brain tissues remains limited. Future targeted proteomic approaches, including sensitive immunoassays like those developed for cryptic HDGFL2^72^, are warranted to confirm and quantify these cryptic products in the brain. Despite these limitations, the integration of human tissue and neuronal models in this study offers a robust framework to dissect the molecular consequences of TDP-43 dysfunction and highlights the potential of CE burden as a mechanistic and biomarker-relevant axis across diverse neurodegenerative diseases.

## Availability of data

Raw mass spectrometry data and pre- and post-processed hippocampus and iNeuron protein abundance data and case metadata related to this manuscript are available at https://www.synapse.org/Synapse:syn65658835.

## Author contributions

Conceptualization, A.N.T., Z.T.M., P.T.S., and N.T.S.; Methodology, A.N.T., A.S., L.P., D.M.D., M.E.W., E.B.D., L.P., P.T.M., Z.T.M., J.D.G and N.T.S; Investigation, A.N.T., M.C. A.S., L.P., Z.T,M., J.D.G., and N.T.S.; Formal Analysis, A.N.T., E.B.D., A.S., N.T.S.; Writing – Original Draft, A.N.T., Z.T.M., and N.T.S.; Writing – Review & Editing, A.N.T., A.S., L.P., D.M.D., L.P., M.E.W., P.T.N., A.I.L., Z.T,M., J.D.G., and N.T.S.; Funding Acquisition, J.D.G., and N.T.S.; Resources, P.T,M., M.E.W., L.P., J.D.G., and N.T.S., and.; Supervision, Z.T.M., J.D.G., and N.T.S. All authors read and approved the final manuscript.

## Acknowledgements

Funding: This study was supported by the following National Institutes of Health funding mechanisms: U01AG061357 (AIL and NTS) and P30AG066511 (AIL), P01NS084974 (LP and JDG), R01NS132330 (LP) and the Foundations for the National Institute of Health AMP-AD 2.0 grant (NTS and AIL). The authors wish to acknowledge the selfless donation of tissue by the patients included in this study. In addition, we would like to thank the Glass and Seyfried laboratory members that offered feedback on early iterations of this project.

## Figure Legends

**Supplemental Figure 1.**
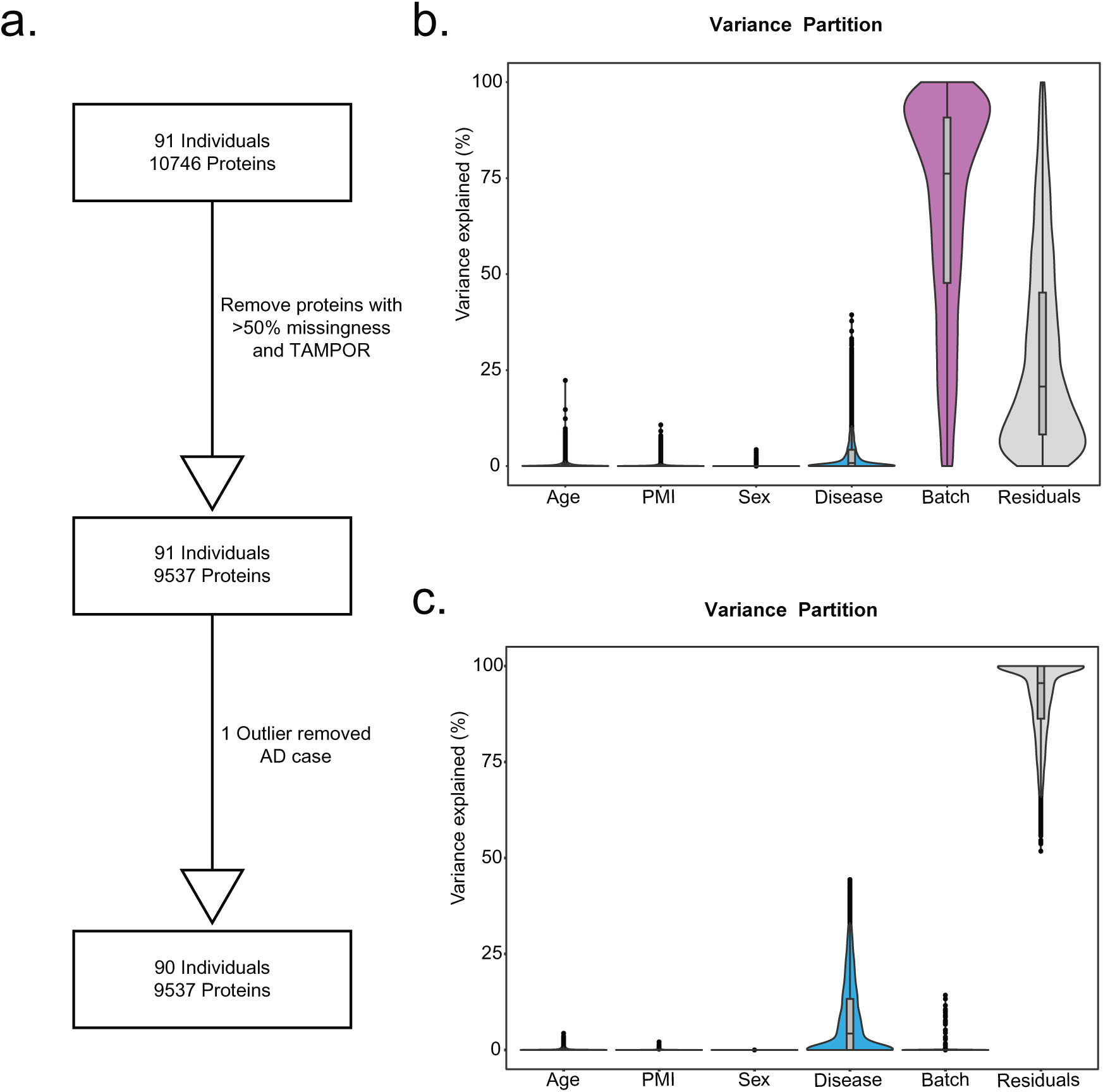
Batch correction and normalization of hippocampus proteomic dataset. A. Experimental design outlining quality control steps, including reductions in sample size and protein count. A variance partition analysis identifies proteins influenced by technical artifacts. B. Median centering of TMT-MS data after TAMPOR processing reduces batch effects. C. Post-regression processing removes batch effects and additionally regresses out age, sex, and postmortem interval (PMI).

**Supplemental Figure 2.**
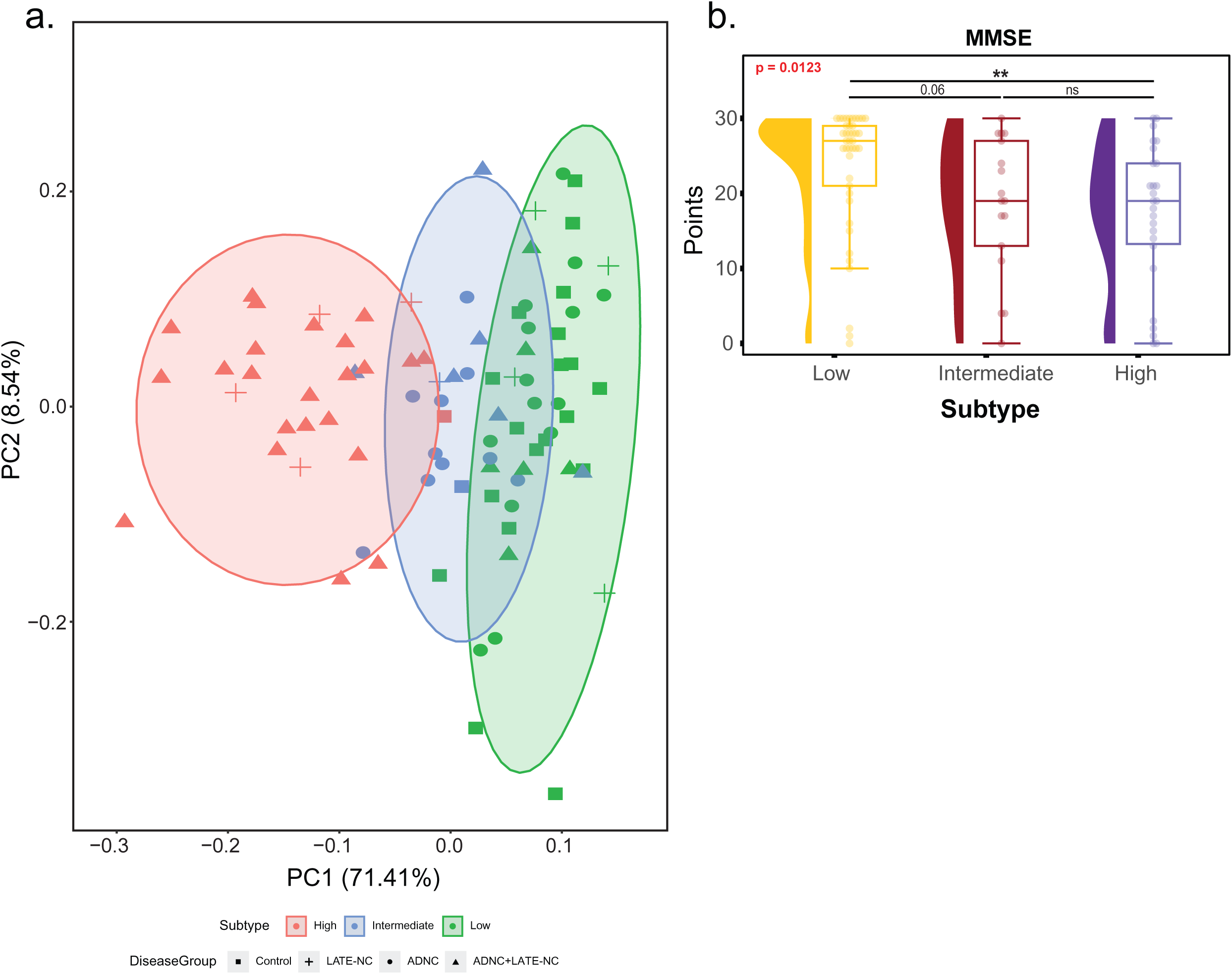
Cryptic exon (CE) profiles resolve molecular subtypes associated with cognition. A. Principal Component Analysis (PCA) shows clear separation between high and low CE burden groups, with intermediates showing mixed profiles. Ellipses represent author added emphasis. B. MMSE scores are compared across subtypes using rainfall plots, including boxplots and score distributions. One-way ANOVA p-values are bolded in red; t-test significance is indicated by asterisks (*p < 0.05, **p < 0.01, ***p < 0.001).

**Supplemental Figure 3.**
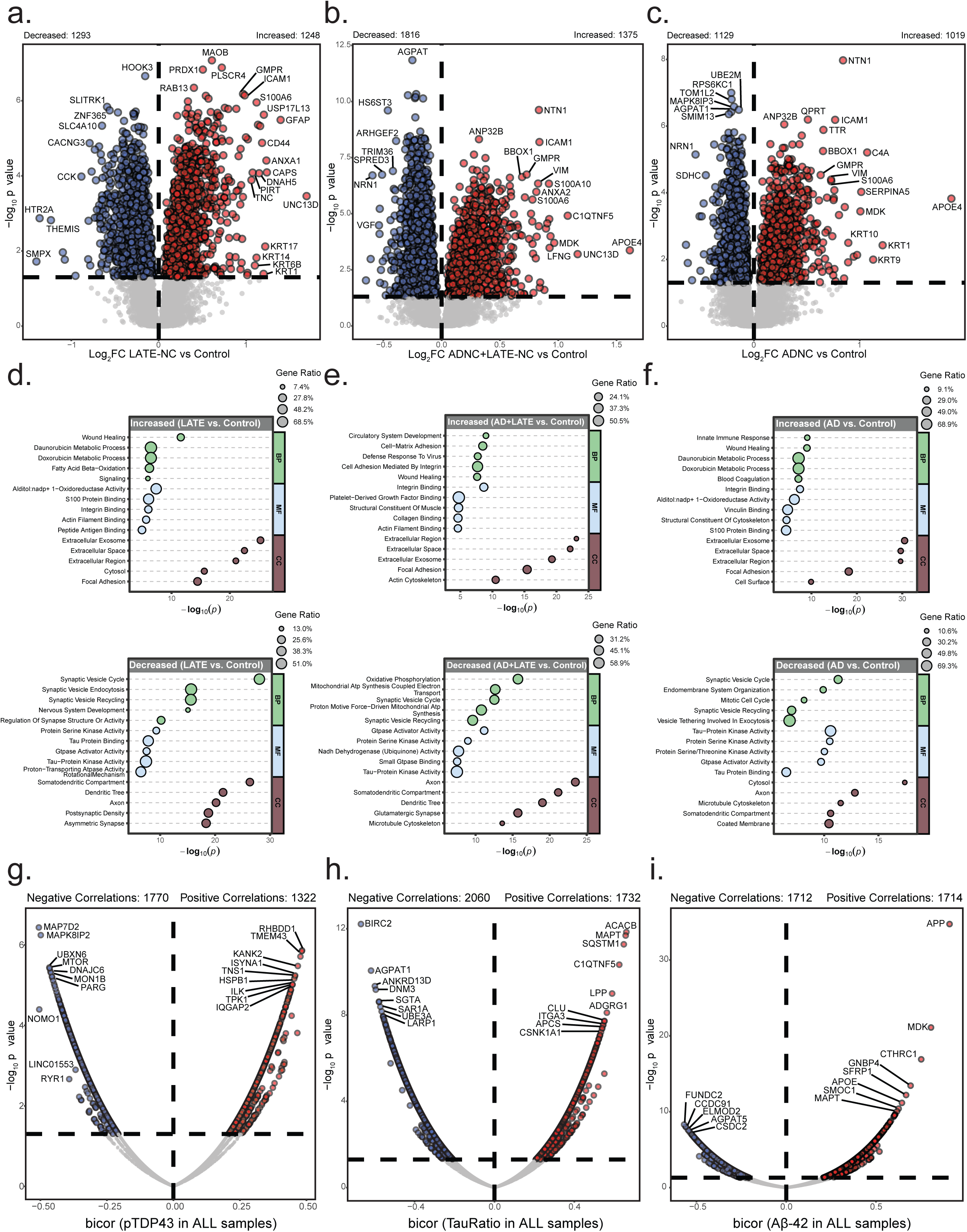
Disease-specific proteomic signatures across clinicopathological subgroups. A–C. Volcano plots of differentially abundant proteins between LATE-NC, ADNC+LATE-NC, ADNC, and control hippocampus samples. Red = increased; blue = decreased; grey = non-significant. D–F. Gene Ontology (GO) enrichment for upregulated (top) and downregulated (bottom) proteins. Circle size represents gene ratio. GO terms categorized by Biological Process (BP), Molecular Function (MF), and Cellular Component (CC). G–I. BiCor correlations between individual proteins and key ADNC/LATE-NC biomarkers (pTDP-43, pTau/Tau, Aβ42). Top correlated proteins are labeled.

**Supplemental Figure 4.**
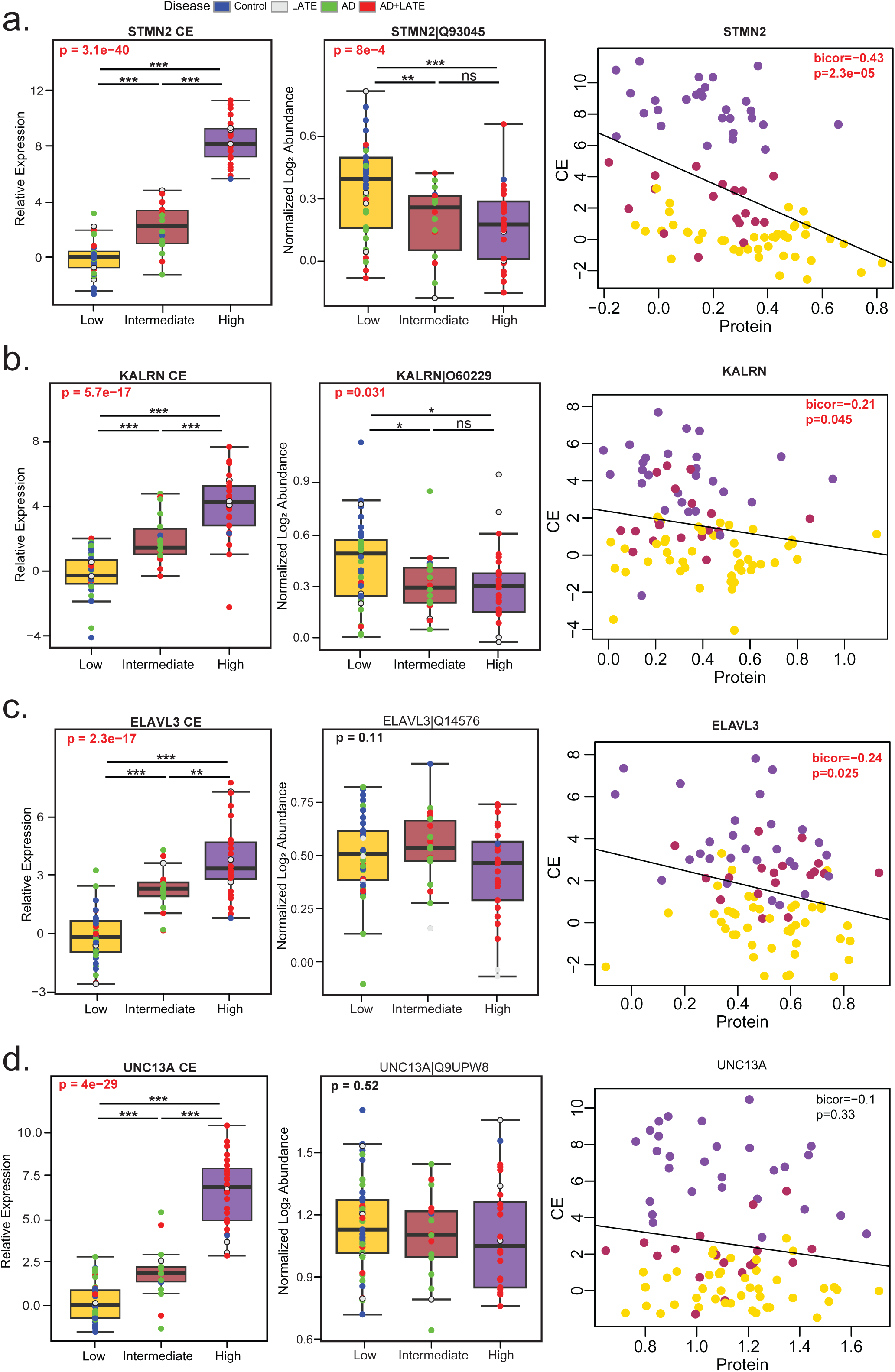
Inverse relationship between cryptic exon (CE) abundance and protein levels. Boxplots and scatterplots show cryptic exon burden, total protein abundance, and their relationship for STMN2, KALRN, ELAVL3, and UNC13A. Points are colored by diagnosis. ANOVA p-values are bold and red where CE burden significantly affects protein/peptide levels. Lines at the top of plots represent t-tests with significance denoted by asterisks (*p < 0.05, **p < 0.01, ***p < 0.001). BiCor and standard correlation p-values are also reported.

**Supplemental Figure 5.**
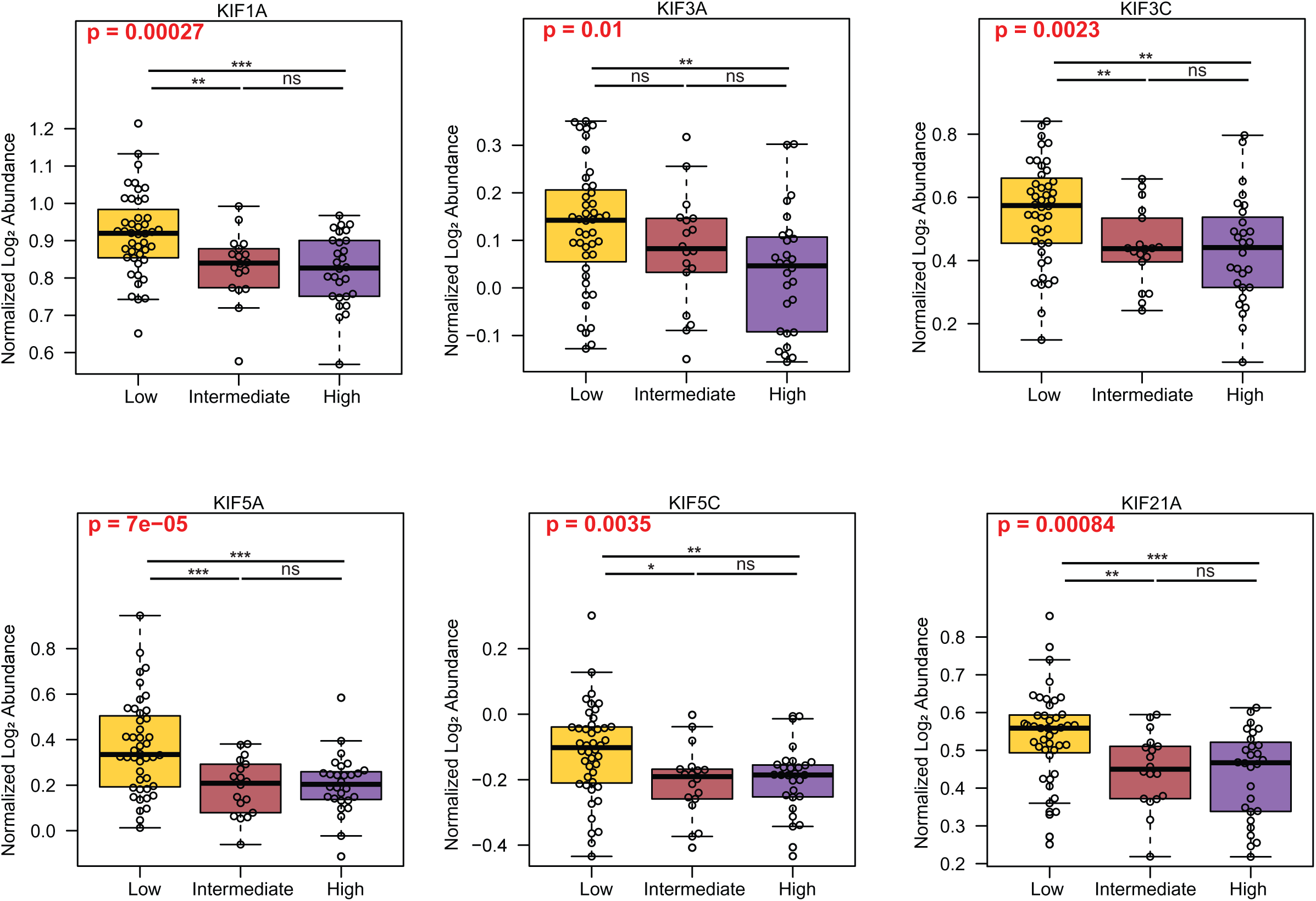
Differential abundance of kinesin proteins across cryptic exon (CE) subtypes. ANOVA results (bold/red) indicate molecular subtype significantly affects kinesin protein abundance. Lines above plots show pairwise t-tests with significance denoted by asterisks (*p < 0.05, **p < 0.01, ***p < 0.001).

**Supplemental Figure 6.**
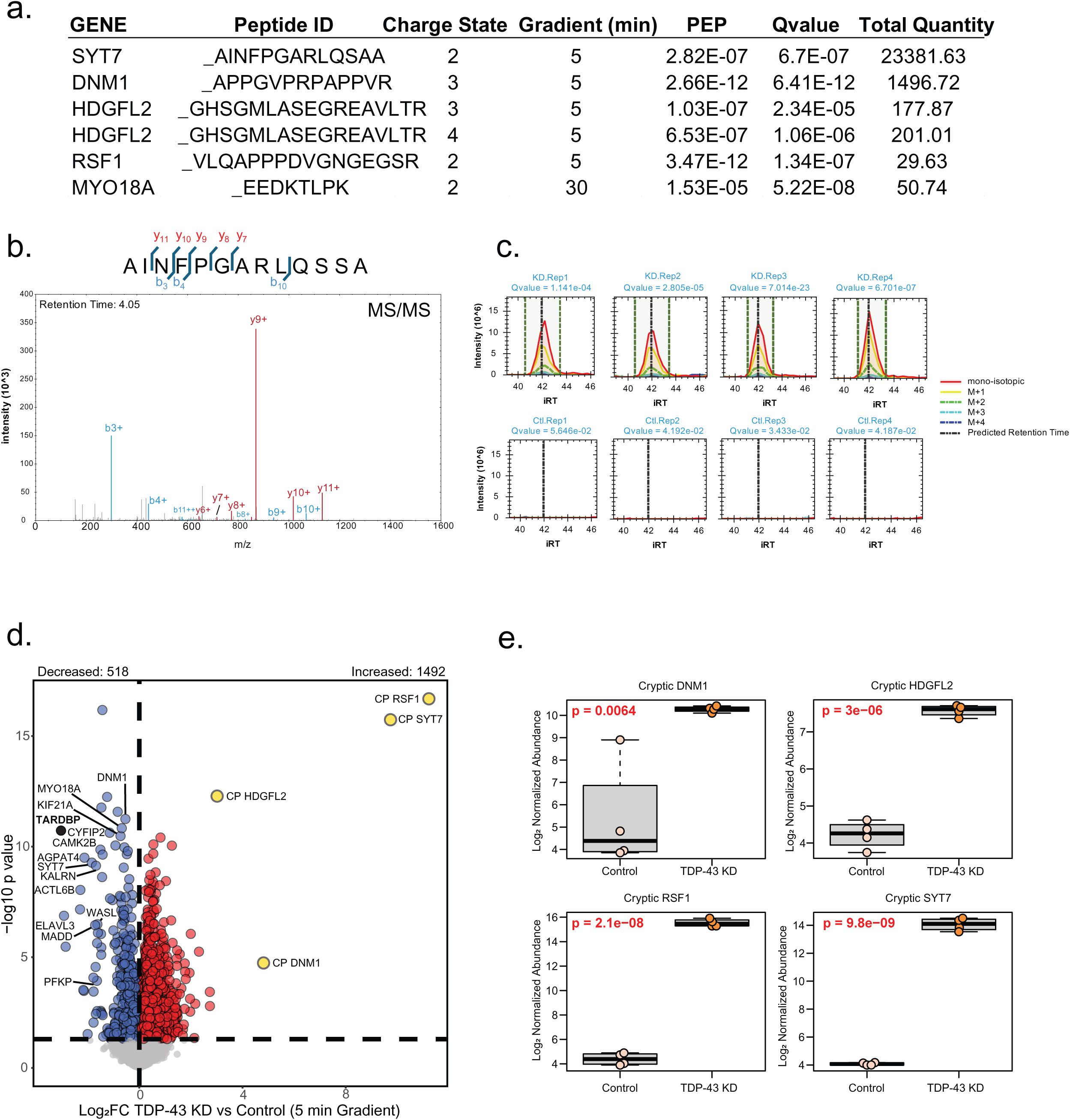
Detection of translated cryptic peptides in TDP-43 KD iNeurons. A. Detected cryptic peptides, including sequence, charge state, and quality metrics. B. MS/MS spectrum identifying a predicted SYT7 cryptic peptide generated by CE inclusion. C. Representative extracted ion chromatogram of MS1 precursor ion intensity (across isotopes) for the SYT7 cryptic peptide was detected in four independent replicates of TDP-43 KD iNeurons, but not control replicates. D. Volcano plot comparing protein abundance between KD and WT in 5-minute LC gradient. Significance color-coded as in Figure 6. E. Individual cryptic peptides show consistent differential abundance between TDP-43 KD and control samples by one-way ANOVA.

